# Reduced glutamate decarboxylase 1 underlies morphine-promoted lung metastasis of triple-negative breast cancer in mice

**DOI:** 10.1101/2023.09.14.557710

**Authors:** Shih-Hong Chen, Chien-Hung Shih, Ting-Ling Ke, Chia-Ni Hsiung, Kuo-Chin Chen, Zi-Xuan Huang, Tsung-Hsien Chuang, Li-Kuei Chen, Linyi Chen

## Abstract

**Introduction:** Morphine is commonly used for cancer-related pain management. Long-term morphine use is not only addictive but also has been associated with risk factor for cancer.

**Methods:** We intraperitoneally administered morphine to mice for 14 days and then implanted EO771 cells, triple negative breast cancer cells, into their mammary fat pad. After primary tumors were removed on 38th day, a subset of mice were continuously giving saline or morphine until the 68th day. Tumor size, organ metastasis, and tumor RNA expression were analyzed.

**Results:** Our results revealed that long-term morphine treatment increased lung metastasis in the triple-negative breast cancer mouse model. To determine cellular pathways responsible for morphine-mediated metastasis, we performed RNA sequencing analysis to compare transcriptional profiles during metastasis. Transcriptional analysis revealed a significant number of genes down-regulated by morphine treatment. Based on pathway analysis, we focused on the novel effect of morphine on down-regulating taurine/hypotaurine biosynthesis. Considering that morphine, droperidol (dopamine receptor antagonist), and naloxone (opioid receptor antagonist) may act through opioid receptor or dopamine receptor, we further demonstrated that taurine reduced EO771 cell invasion caused by morphine, but not by droperidol, or naloxone treatment. In addition, morphine treatment significantly reduced the expression of *GAD1*, one of the enzymes required for biosynthesis of taurine, whereas droperidol and naloxone did not.

**Conclusion:** These novel findings of morphine reduces *GAD1* level and taurine reverses invasion suggest that taurine could potentially be employed as a supplement for triple negative breast cancer patients using morphine as pain management.

**Key Messages:**

- Morphine usage has been implicated in modulating immune system and affecting cancer progression.
- Long-term usage of morphine promotes metastasis of triple negative breast cancer through reducing taurine biosynthesis.
- These novel findings of morphine reduces GAD1 level and taurine reverses invasion suggest that taurine could potentially be employed as a supplement for triple negative breast cancer patients using morphine as pain management.

## Introduction

Breast cancer has been a great concern in clinical practice, consisting a distressing 39% of newly diagnosed cancer cases and accounting for 44% of deaths in Asian countries. Triple-negative breast cancer is characterized by low expression levels of the estrogen receptor, progesterone receptor, and human epidermal growth factor receptor. The rate of metastasis for triple-negative breast cancer is the highest compared to other breast cancer types, and the survival probability of triple-negative breast cancer patients is thus lower due to the lack of a therapeutic target.

Breast cancer and metastasis-related pain syndromes pose significant challenges during clinical treatment, as over 70% of breast cancer patients encounter symptoms associated with pain[1]. Morphine is commonly used for patients with chronic pain syndrome, including cancer related-pain syndrome, may regularly take morphine. Chronic usage of >120 mg/day in clinical settings may lead to opioid tolerance[2], and morphine dosage needs to be increased as the progression of pain due to tolerance[3]. While morphine exposure itself may not increase the risk of breast cancer, it can potentially worsen the survival outcomes of breast cancer patients. Recent studies have reported the adverse effect of morphine on tumor growth and metastasis, including in triple-negative breast cancer[4, 5]. Mechanisms underlying morphine-mediated tumor growth and metastasis remain unclear. For example, Mathew et al. reported that knockout of the μ-opioid receptor (MOR) or the use of a MOR antagonist considerably inhibited tumor growth[6]. Nonetheless, clinical data point to a greater potential of morphine in promoting metastasis rather than inhibiting it[7]. To our knowledge, only few studies reported the critical importance of advancing medical practice in addressing breast cancer, metastasis, and morphine usage.

The present study compared the gene expression profiles of metastatic tumors caused by long-term morphine exposure and spontaneous triple-negative breast tumors. Our data indicated that long-term morphine administration increased the progression and metastasis of triple negative breast cancer cell-implanted mice. These findings urge the importance of optimal management of morphine usage in effectively tackling breast cancer and metastasis.

## Materials and Methods

### Reagents

RPMI-1640, fetal bovine serum (FBS), and 6-diamidino-2-phenylindole were purchased from Invitrogen (Carlsbad, CA, USA). Additionally, 3-(4,5 dimethylthiazol-2-tl)-2,3-diphenyltetrazolium bromide (MTT) was purchased from Sigma-Aldrich (STL, USA). Bovine serum albumin (BSA) was purchased from Santa Cruz Biotechnology (SantaCruz, CA, USA). The SPLInser polyethylene terephthalate membrane (pore size: 8.0 μm) was obtained from SPL Lifesciences (Pocheon, South Korea). Matrigel was purchased from BD Biosciences (San Jose, CA, USA). TRIzol reagent was purchased from Invitrogen. The SYBR green master mix, reverse transcription kit, and StepOnePlus PCR system were purchased from Applied Biosystems (Foster City, CA, USA). Taurine was from Sigma-Aldrich (Saint Louis, MO, USA), Morphine sulfate was from Taiwan Food and Drug Administration, Ministry of Health and Welfare, Taiwan. Droperidol was purchased from the Excelsior Pharmatech Labs (Taipei, Taiwan) and Naloxone was from UniPharma (Taipei, Taiwan).

### Cell culture, animal handling, and ethics statement

EO771 cells [8–10] (obtained from Dr. Tsung-Hsien Chuang, National Health Research Institutes, Taiwan) were maintained in six-well plates with RPMI-1640 supplemented with 10% FBS at 37°C and 10% CO_2_. Female, 12–14-week-old C57BL/6 mice were purchased from the National Laboratory Animal Center, Taiwan, and housed in a 12/12-hour light/dark cycle.

Experimental mice (n = 11, C57BL/6Jarl, female) were intraperitoneally injected with morphine (10 mg/kg/day) or equal volume of saline for the indicated days (Figure 2) randomly, and the EO771 cells were implanted into the mammary fat pad (2 × 10^5^ cells in 150 μL of PBS) on the 14th day. The raising conditions and cage placements were identical in both groups according to NTHU animal room protocol. Possible side effects of morphine were monitored, including low appetite, no fecal passage, and else. To evaluate the antinociceptive response of mice, we employed the tail-flick assay [11]. Prior to the administration of morphine, a baseline measurement was obtained. Subsequently, repeated measurements were taken at two time points: 30 minutes after the injection of morphine on the 14th and 37th day. To prevent tissue damage, a cut-point of 10 seconds was established as a threshold. Specifically, a latency delay of more than 3 seconds was considered indicative of the effectiveness of morphine in producing antinociception[12]. Tumor size was measured every 3 days from the 31st day. Tumor volume calculations were obtained using the formula Volume = (Width^2^ × Length)/2 for caliper measurements. The measurements were conducted simultaneously and on the same working platform for both the experimental and control groups. Thirty-eight days after the first morphine injection, the tumor was resected under general anesthesia (1.5% isoflurane with oxygen) in National Tsing Hua University (NTHU) animal care facility. After removing the primary tumors, six of the saline-treated group and four of the morphine-treated group were continuously given saline or morphine and euthanized on the 68th day. Their liver and lung were also harvested to evaluate metastasis on day 68. All animal experiments were conducted in accordance with the guidelines of the Laboratory Animal Center of NTHU, Taiwan. Animal usage protocols were reviewed and approved by NTHU’s Institutional Animal Care and Use Committee (approval number: 10607). The assistant personnel responsible for the care, measurements, and sacrifices do not participate in the subsequent data analysis.

### Cell proliferation and invasion assays

EO771 cells used for proliferation and invasion assays were the same source and passage numbers. Triplicates were used for each independent experiment. At least three independent experiments were performed. To determine the effect of morphine on cell viability/proliferation, EO771 cells cultured in a 96-well plate were treated with different morphine concentrations (1, 10, or 100 μM) for 48 h. After treatment, 3-(4,5-dimethylthiazol-2-yl)-2,5-diphenyltetrazolium bromide (MTT) were added to each well at final concentration 0.5 mg/ml, and then incubated for 3 h at 37°C in dark. The supernatants were removed and MTT product formazan at the bottom of plates was dissolved by 100 μl 100% DMSO. Absorbance of formazan was measured at 550 nm with a microplate reader. Subsequently, the absorbance of optical density (OD) at a wavelength of 565nm was measured. The percentage of cell viability was determined using the following equation:

Viability (%) = 100 x OD (mean value of test)/ OD (mean value of negative control) Morphine stock solution was serial-diluted with RPMI medium (without FBS), thus RPMI was added to the culture as the negative control (0 morphine).

To determine cell invasion, the Boyden chamber assays were performed to analyze cell invasion ability. A transparent polyethylene terephthalate (PET) membrane (8 μm) insert was coated with Matrigel in 24-well plates, EO771 cells were suspended in serum-free RPMI medium (containing 0.1% w/v BSA and different concentrations of morphine, naloxone, droperidol, taurine) and 2 x 10^5^ cells were plated onto solidified Matrigel. RPMI medium (supplemented with 10% FBS and different concentrations of morphine, naloxone, droperidol, taurine) was added to the lower chamber as a chemoattractant to attract cell invasion. After incubation for 26 h, cells migrated to the bottom side of PET membrane were fixed with 4% paraformaldehyde and stained with crystal violet. Images of crystal violet-stained cells were taken using Zeiss Observer Z1 microscope (Zeiss, Jena, Germany). Number of cells was counted using Image J software (https://imagej.nih.gov/ij/index.html). Relative invasion ability was calculated as cell number per chamber and normalized to “0 morphine” control (=1).

### Real-time quantitative PCR

RNAs of EO771 cells were collected using TRIzol and mRNAs (2 μg) were reverse transcribed to cDNA. Real-time PCR with SYBR Green detection was performed using an ABI PRISM 7500 sequence detection system (Applied Biosystems). Glyceraldehyde-3-phosphate dehydrogenase (GAPDH) was used as a control. Primers were listed below: GAD1_F: 5’-GCG GGA GCG GAT CCT AAT A-3’; GAD1_R: 5’-TGG TGC ATC CAT GGG CTA C-3’; CDO1_F: 5’-GAG GGA AAA CCA GTG TGC CTA C-3’; CDO1_R: 5’-CCT GTT CTC TGG TCA AAG GCG T-3’; GAPDH_F: 5’-ATG TTT GTG ATG GGT GTG AA-3’; GAPDH_R: 5’-ATG CCA AAG TTG TCA TGG AT-3’.

### RNA-sequencing and data analysis

Total RNAs were extracted from collected tumors with a High-Capacity cDNA Reverse Transcription Kit (Applied Biosystems). Agilent 2100 Bioanalyzer–with Agilent RNA 6000 Nano kit was used to determine the quality of RNAs followed by the RNA sequencing using Illumina HiSeq 4000, conducted by Genomics, a biotech company in Taiwan (https://www.genomics.com.tw/). According to standard protocol as following:

#### Library preparation and sequencing

The purified RNA was used for the preparation of the sequencing library by TruSeq Stranded mRNA Library Prep Kit (Illumina, San Diego, CA, USA) following the manufacturer’s recommendations. Briefly, mRNA was purified from total RNA (1 μg) by oligo(dT)-coupled magnetic beads and fragmented into small pieces under elevated temperature. The first-strand cDNA was synthesized using reverse transcriptase and random primers. After the generation of double-strand cDNA and adenylation on 3’ ends of DNA fragments, the adaptors were ligated and purified with AMPure XP system (Beckman Coulter, Beverly, USA). The quality of the libraries was assessed on the Agilent Bioanalyzer 2100 system and a Real-Time PCR system. The qualified libraries were then sequenced on an Illumina NovaSeq 6000 platform with 150 bp paired-end reads generated by Genomics, BioSci & Tech Co., New Taipei City, Taiwan.

#### Bioinformatics Analysis

The RNA-seq was sequencing by Illumina NovaSeq 6000. Low quality bases (< Q20) and adapters were removed by “Trimmomatic v0.36”. Whole Transcriptome was de-novo assembled by “Trinity v2.8.4”, then using “cd-hit-est v4.7” to generate uni-transcript with 95% clustering cutoff. Each sample paired-end reads were aligned to transcript fasta using “Bowtie2 v2.3.4.3”, calculating gene expression count by “RSEM v1.2.28”. Differentially expressed genes (DEGs) were calculated by “egdeR v3.24.1”. Transcripts were downstreaming with ORF prediction by “Transdecoder v5.3.0” and the functional analysis, include signalP, TMHMM, PFAM, enzyme detection, COG, GO analysis, were using “Trinotate v3.1.1”. Transcript nucleotide fasta was annotated by “BLASTX v2.5.0”, and predicted amino acid fasta was annotated by “BLASTP v2.5.0”. KEGG pathway graph is coverted from EC number by ec2kegg.pl, which is a perl script contributed by Aleksey Porollo.

Purified RNA was used for preparation of the sequencing library by using the TruSeq Stranded mRNA Library Prep Kit (Illumina, San Diego, CA, USA) in accordance with the manufacturer’s instructions. Briefly, mRNA was purified from total RNA (1 μg) by using oligo (dT)-coupled magnetic beads and fragmented into small pieces under elevated temperature. First-strand cDNA was synthesized using reverse transcriptase and random primers. After the generation of double-stranded cDNA and adenylation on the 3′-end of DNA fragments, adaptors were ligated and purified using the AMPure XP system (Beckman Coulter, Beverly, USA). The quality of the libraries was examined using the Agilent Bioanalyzer 2100 system and a real-time polymerase chain reaction system. The qualified libraries were sequenced on an Illumina NovaSeq 6000 platform with 150-bp paired-end reads generated by Genomics (BioSci & Tech Co., New Taipei City, Taiwan). All experiments were performed in Genomics Core through the Genomics Core laboratory process (https://en.genomics.com.tw/).

The low-quality bases and sequences from adapters in raw data were removed using Trimmomatic (version 0.39) [13]. The filtered reads were aligned to reference genomes by using Bowtie2 (version 2.3.4.1) [14]. The user-friendly software RSEM (version 1.2.28) was used for quantification of transcript abundance [15]. Normalization and identification of differentially expressed genes (DEGs) were carried out by using the R package edgeR. We applied a generalized linear model to estimated effect sizes and p-values of differentially expressed genes between morphine and saline treatments [16, 17]. The functional enrichment analysis of Gene Ontology (GO) terms and Kyoto Encyclopedia of Genes and Genomes (KEGG) pathways among gene clusters was performed using an R package called ClusterProfiler (version 3.6.0) [18–20] and analyzed using an online bioinformatics tool (Database for Annotation, Visualization, and Integrated Discovery, DAVID; https://david.ncifcrf.gov) [21]. Principal component analysis (PCA) and volcano plot generation were performed using the R package ggplot2. A heatmap was generated using the R package heatmap.3.

To analyze protein–protein interactions (PPIs), we used STRING (https://string-db.org) [22]. We employed the Cytoscape plugin Cytohubba to determine the highest scores of hub genes [23]. Additional pathway analysis was performed using the Cytoscape plugin ClueGo [24]. Transcription factors were analyzed using TRRUST (https://www.grnpedia.org/trrust/).

To determine the relationship between the selected hub genes or pathways and survival, we compared the identified hub genes and pathways by using data from The Cancer Genome Atlas (TCGA) database. The UCSC Xena website (https://xenabrowser.net) is an online database used for determining the correlations among genotypes, phenotypes, and survival in TCGA breast cancer dataset. We applied hub and transcription genes in UCSC Xena to determine the association of overall survival with gene expression. Patients were divided into high and low gene expression groups for the survival analysis. For the pathway analysis, we determined the total number of genes involved in each pathway and analyzed the relationship between relapse-free survival and pathway expression [25].

### Statistical Analysis

Difference between un-treated and different morphine dose-treated samples for MTT and invasion assays was determined by one-way ANOVA. For multiple comparison, we adjusted P value by bonfernonni method. Each morphine-treated sample was normalized to untreated sample (=1).

We evaluated the difference of tumor volumes between saline- and morphine-treated groups for 17, 20, 23 days, we excluded outliers which were outside mean±3 standard deviation, and used two-way ANOVA and Tukey post-hoc test. Relative invasion was performed using Boyden chamber assays and analyzed using Student’s t-test. For RNA-seq analysis, normalization and identification of differentially expressed genes (DEGs) were carried out by using the R package edgeR. We applied a generalized linear model to estimate effect sizes and FDR p-values of differentially expressed genes between morphine and saline treatments. For the survival analysis, we used Kaplan-Meier and log-rank test. All the statistical methods were used by R (version 4.0.2).

## Results

### Effect of morphine on cell viability, invasion, tumor growth, and metastasis

Triple-negative breast cancer is highly metastatic and has worse prognosis than other breast cancer subtypes [26]. The EO771 cell line is derived from the C57 BL/6 mouse model of spontaneous breast cancer, and this is an established model for studying triple-negative breast cancer [27]. To investigate the effect of long-term morphine usage on breast cancer cells, EO771 cells were incubated with morphine at the concentrations of 0, 1, 10, or 100 μM for 48 h. Our results revealed that up to 100 μM morphine did not affect cell proliferation compared with the mock-treated control (Figure 1). The Boyden chamber assays were performed to examine the effect of morphine on the invasion ability of the EO771 cells within 26 h. The EO771 cells treated with ≥1 μM morphine exhibited at least twice the invasion ability (Figure 1) of the control group. These results suggest that morphine significantly enhances the invasion ability of EO771 cells but does not affect their proliferation.

**Figure 1.**
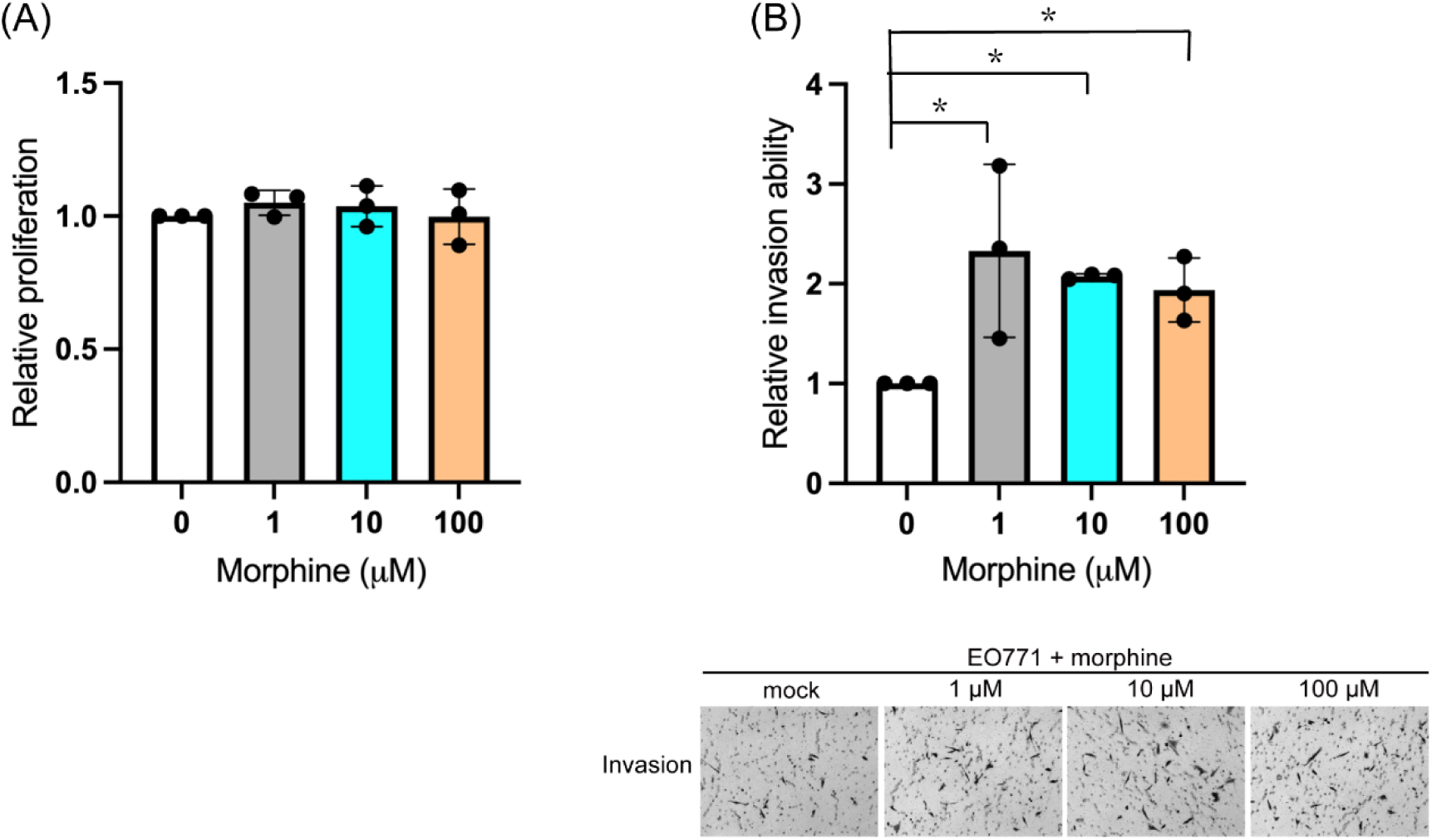
Effect of morphine on the proliferation and invasion of EO771 cells. (A) EO771 cells were treated with 0–100 μM morphine. The MTT assays were performed to determine relative cell proliferation/viability after 48 h of treatment. (B) EO771 cells were treated with 0–100 μM morphine for 26 h, and the Boyden chamber assays were performed to determine relative invasion ability (* indicates a significant difference of P < 0.05 between 0 and morphine-treated groups, determined using the Student’s t test.). Images of invaded cells are shown. Data were from three independent experiments.

While morphine is used for pain management, this study focuses on the effect of long-term use of morphine on cancer progression, thus no additional pain-initiating agent or pain-relief medicine were used to avoid drug-drug interaction. To mimic the effects of long-term morphine usage in humans, the mice were intraperitoneally administered 10 mg/kg morphine or same volume of saline daily for 2 weeks before EO771 cell implantation to fat pad. The tail-flick test was used to ensure delayed nociception to a heat stimulus for morphine-treated mice. Figure 2A presents a schematic of the experimental approach including morphine injection and tumor measurement. Thirty-one days after the first morphine injection and 17 days after implantation of the EO771 cells, tumor size was measured every 3 days. T Morphine pre-treatment resulted in a significant increase in the tumor volume. After removing primary tumors on day 38, 6 saline-treated mice and 4 morphine-treated mice were followed-up for potential metastasis till day 68. On the 54th day after tumor cell implantation, lung metastasis developed in 3 out of the 6 mice in the saline group and in all 4 mice in the morphine-treated group. The tumor volume was measured on the 17^th^, 20^th^, and 23^rd^ day after tumor cell implantation. For data analysis, we excluded outliers which were outside mean±3 standard deviation. For saline group, there were 10, 10, and 8 mice on the 17^th^, 20^th^, and 23^rd^ day, respectively; for morphine groupthere were 9, 9, and 9 mice on the 17^th^, 20^th^, and 23^rd^ day, respectively. The tumor sizes were compared among days and treatments using two-way ANOVA and Tukey for post-hoc test, Tumor volumes on the 17^th^ day and the 20^th^ day were not significant difference. On the 23^rd^ day, tumors derived from morphine-treated groups were significantly larger than those from saline-treated groups (P=0.0001). When comparing among different treatments and time periods, only tumors from the 23^rd^ day were significantly different from those from 17^th^ and 20^th^ (23^rd^ vs 17^th^ P=0.00035, 23^rd^ vs 20^th^ P=0.0022). Tumors among saline-treated groups were not significantly different (Figure 2B). The incidence of lung metastasis was higher in the morphine-treated group compared to saline-treated group (Figure 2C). The average total area of tumors in lung, on the other hand, was increased in morphine-treated mice, but without significant difference (Figure 2C-D). The total area of tumor growth in lung represents colonization of metastasized tumors. The EO771-implanted mice exhibited heterogeneity in terms of small or large tumors, thus we categorized saline control groups into S1–S3 (Table S1). To elucidate the mechanism through which morphine affects tumor formation and metastasis, we extracted total RNA from isolated tumors for RNA sequencing analysis. DEGs were compared between the morphine and control groups. For the controls, RNA samples extracted from tumors with size larger than 50% of the mean volume without metastasis, tumors with size less than 50% of the mean volume without metastasis, and tumors with metastasis were designated the Saline 1 (S1), Saline 2 (S2), and Saline 3 (S3) groups, respectively (Table S1). In the morphine-treated groups, RNA samples extracted from tumors from mice that were not followed-up for metastasis --- morphine 1 (M1), and those that were followed-up metastasis --- morphine 2 (M2) (morphine+metastasis) groups, respectively. In terms of gene expression profiles, PCA indicated that the first (PC1) and second (PC2) principal components represented 42.5% and 28.2% of all variables, respectively, and explained 70.7% of the total variance. The S1 and S2 groups were located in the upper-right quadrant. The M1 group was located in the lower-right quadrant. The S3 and M2 groups were located in the upper-left and lower-left quadrant, respectively. This PCA analysis revealed distinct transcriptional profiles among groups, with clear distinction depending on morphine treatment or metastasis (Figure S1A). A heatmap was plotted to examine the correlations of gene expression profiles among S1, S2, S3, M1 and M2. Clusters of S1, S3 were different from M1 and M2 whereas S2 was different from other samples. This analysis suggests that the mechanism underlies the tumor growth and spontaneous metastasis of triple negative breast is likely different from that of promoted by morphine (Figure S1B).

**Figure 2.**
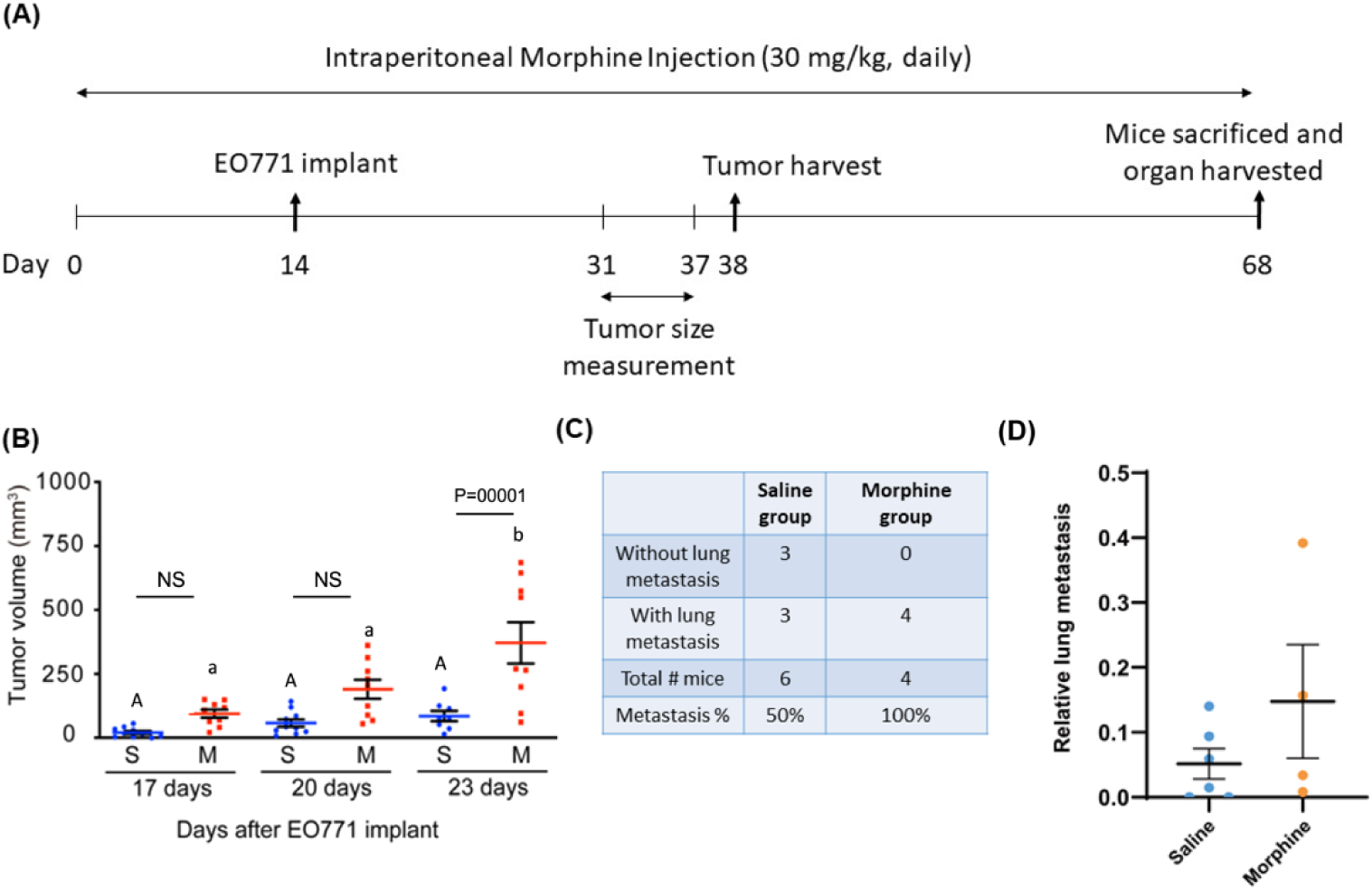
Analysis of tumor sizes and metastasis with or without morphine treatment. (A) Mice received an intraperitoneal morphine injection for 14 days. On day 14, EO771 cells were implanted into a fat pad. Tumor size was measured from day 31 to day 37. Primary tumors were collected on day 38; this was followed by suturing the wound area. 6 saline-treated and 4 morphine-treated mice were continuously administered saline or morphine till day 68. Liver and lung samples were collected on day 68 to determine possible metastasis. (B) Tumor volume (mm^3^) was measured on the 17^th^, 20^th^, and 23^r^d day after EO771 implantation. The original animal numbers were 11 in saline group and 10 in morphine group. For data analysis, we excluded outliers which were outside mean±3 standard deviation. In saline group, n= 10, 10, 8 on the 17^th^, 20^th^, and 23^rd^ day, respectively; for morphine group n= 9, 9, 9 on the 17^th^, 20^th^, and 23^r^d day, respectively. The A, a, and b showed statistical results of different saline and morphine treatment days (17^th^, 20^th^ and 23^rd^) using two-way ANOVA and Tukey post-hoc test. The “A” and “a” showed no difference in these three conditions by day. The “a” and “b” showed that morphine treatment group from the 23 days was higher than those of 17 and 20 days. (C) Metastasis % was calculated based on the number of mice with lung metastasis in saline-treated and morphine-treated mice. (D) Relative area of tumors in lung for saline-treated and morphine-treated mice. Student’s t test was used for statistical analysis, P=0.357.

### Screening of DEGs

PCA and heatmap analysis demonstrated cluster characteristics and similar gene expression between the S1 and S2 groups. We calculated the average expected gene expression of the S1 and S2 groups and compared the expected gene expression of the M2 group with the average expression of the S1 and S2 groups. In the RNA seq analysis, 12,592 DEGs were identified, of which 401 were identified as dominant genes (false discovery rate [FDR] < 0.05). Among those 401 genes, 32 hub genes were up-regulated (Log_2_ FC >1) and 225 hub genes were down-regulated (Log_2_ FC < −1). The volcano plot illustrated the distribution of up-regulated and down-regulated genes. We observed significantly down-regulated expression in the morphine group which was shown in volcano plot (Figure S2). We used the DAVID tool for performing the DEG enrichment analysis, including GO term and KEGG analyses. We analyzed 12,592 DEGs and identified 32 up-regulated (Log_2_ FC >1) and 225 down-regulated hub genes. We determined the GO terms and KEGG pathways of the identified 257 genes by using DAVID website tool. The findings indicate that the metastatic effect of morphine significantly down-regulates the expression of a subset of genes. We compared morphine influence of gene expression (M2) with saline group (S3) in metastasis tumors, and 12,586 genes were identified. There were 93 dominant up-regulation hub genes and 63 down-regulation hub genes, and volcano plot showed less dominant genes (Figure S2B).

### Functional and pathway enrichment analysis

We used the DAVID tool for performing the DEG enrichment analysis, including GO term and KEGG analyses. We analyzed 12,592 DEGs and identified 32 up-regulated (Log_2_ FC >1) and 225 down-regulated hub genes. We determined the GO terms and KEGG pathways of the identified 257 genes by using DAVID. In the GO analysis, enrichment processes were divided into biological process (BP), molecular function (MF), and cellular compartment (CC) categories. Among the up-regulated 32 genes, 16.1% of the genes were involved in each of the cytosis (GO:0019835), protein processing (GO:0016485), immune responses (GO:0006955), and proteolysis (GO:0006508), respectively. In terms of MFs, 16.1% of the genes were involved in each of serine-type peptidase activity (GO:0008236), serine-type endopeptidase activity (GO:0004252) and peptidase activity (GO:0008233), whereas 19.4% of the genes were involved in hydrolase activity (GO:0016787). In terms of CCs, we identified only the intracellular membrane-bounded organelle pathway (GO:0043231, 19.4%; Table S2). The 225 down-regulated genes were involved in 64 biological processes, of which 20 and 25 were in MF and CC processes (Table S3). The three most common processes in each group were as follows: immune response (GO:0006955, 6.7%,), inflammatory response (GO:0006954, 6.7%), and lipid metabolic process (GO:0006629, 6.3%) in BPs; hydrolase activity (GO:0016787, 12.1%), calcium ion binding (GO:0005509, 10.7%), and protein homodimerization activity (GO:0042803, 8.3%) in MFs; and membranes (GO:0016020, 41.5%), extracellular exosomes (GO:0070062, 30.8%), and extracellular regions (GO:0005576, 25%) in CCs. The results of the KEGG analysis are presented in Table S4. The up-regulated genes belonged to the Legionellosis (mmu05134, 6.5%) pathway. Pathways that were significantly affected by the down-regulated genes included the cytokine–cytokine receptor interaction (mmu04060, count 8), lipolysis regulation in adipocytes (mmu04923, count 7), and cGMP-PKG signaling pathway (mmu04022, count 7). Because few genes were determined to be up-regulated during morphine-promoted metastasis, fewer pathways were identified; they included those involved in proliferation and invasion. The down-regulated genes participated in immune-related responses. Suppressed immune response has been implicated in tumor formation. Thus, morphine usage can be associated with immune responses and tumor formation. Consistent with this evidence, morphine usage can also be associated with metastasis. The expression of genes involved in regulation of the extracellular matrix was observed to be decreased in this study, supporting the aforementioned finding. The genes determined to be involved in lipid metabolism, protein/peptide metabolism, and signal transduction in the KEGG analysis were compatible with those identified in the GO term analysis.

The KEGG analysis demonstrated the dominant down-regulation of arachidonic acid, lipolysis, the renin–angiotensin system, and taurine/hypotaurine metabolism. Down-regulation of these pathways may contribute to tumor invasion. We used the Cytoscape plugin ClueGo to integrate the functional pathways identified in the GO term and KEGG analyses, and the dominant KEGG pathways and top 20 GO terms are illustrated in Figure 3A. Because we identified a considerable number of down-regulated genes, 225 significantly down-regulated genes were selected for the subsequent analysis (FDR < 0.05). The results of the DAVID analysis revealed 10 down-regulated pathways after long-term morphine treatment. Figure 3B presents the relative expression of genes (S1 and S2 average counts, S3 counts, and M2 counts) within each pathway. The GO term analysis indicated the dominant down-regulation of pathways by morphine treatment. Of these, one novel finding is that morphine affects taurine/hypotaurine metabolism which includes three taurine biosynthesis enzymes.

**Figure 3.**
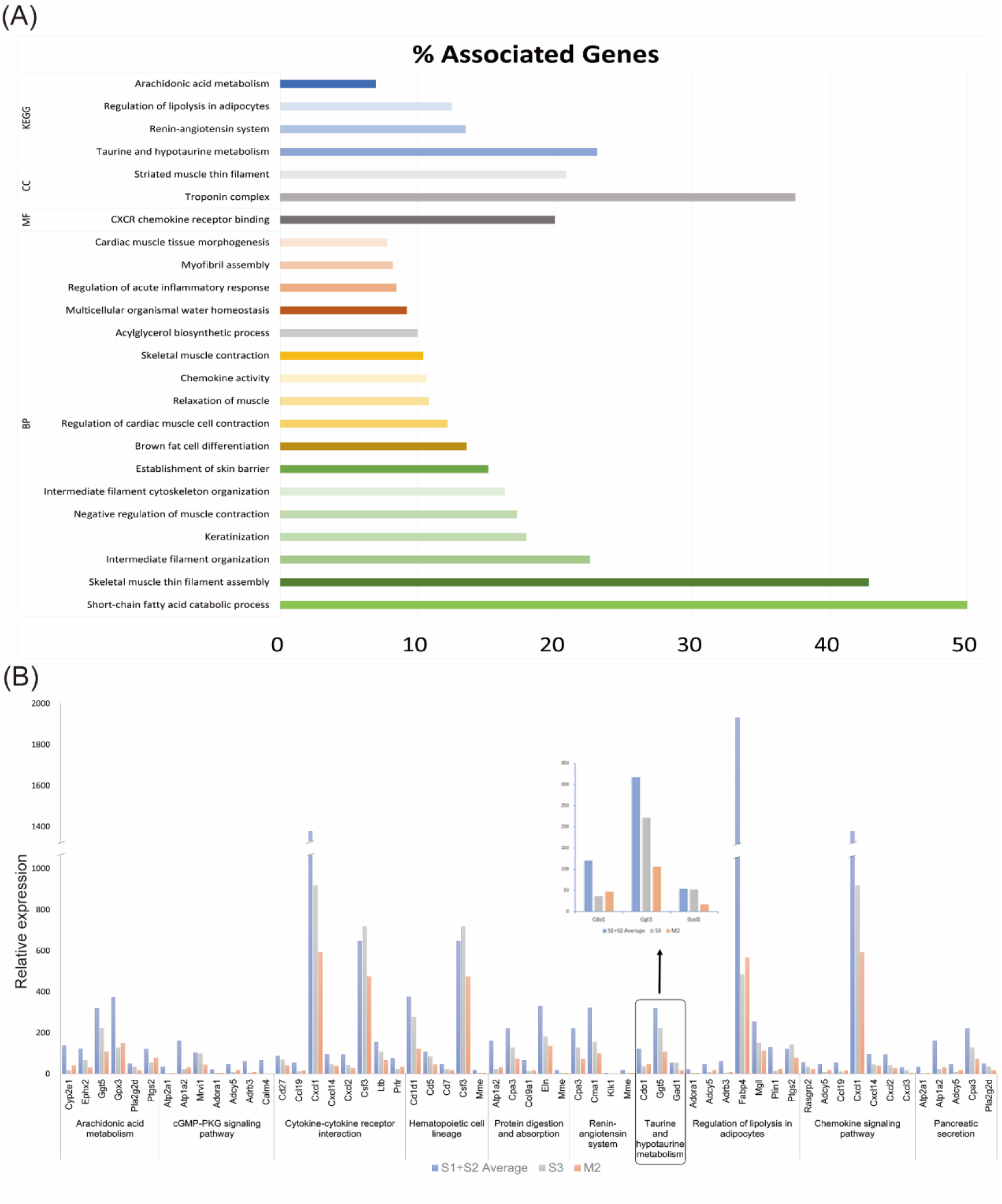
ClueGo analysis to identify significant KEGG pathways and the gene expressions of down-regulated KEGG pathways. (A) The dominant KEGG pathways and top 20 GO terms are illustrated. Relative expression of 10 down-regulated KEGG pathways analyzed using the Cytoscape plugin ClueGo. (B) Average expression counts of the S1 plus S2 groups (blue), S3 group (grey) and M2 group (orange) are compared.

Morphine long-term morphine usage modulates the tumor microenvironment and, in turn, causes these pathways to promote cancer progression and metastasis. We explored the possibility of interaction among these pathways. PPIs of DEGs were determined using the STRING database (Figure S3). To determine the protein count and interaction, we designated a protein as a node and an edge as the interaction between two proteins. Fewer interactions (seven nodes and five edges) were noted among the up-regulated proteins due to the identification of fewer up-regulated genes. We observed more interactions among the down-regulated proteins (156 nodes and 587 edges). A total of 11 down-regulated hub genes (*TTN*, *TCAP*, *ACTA1*, *MYH1*, *TNNT3*, *MYL1*, *MYH4*, *TNNI2*, *ATP2a1*, and *TNNC2*) were selected with the highest total score as indicated by the Cytoscape plugin Cytohubba.

To verify the hub genes and pathways and to determine the relationship between down-regulated hub genes or pathways and patient survival, we examined the candidate genes by using TCGA Breast Cancer (BRCA) datasets (n = 1097). The BRCA dataset is a cohort database and includes information on variations in gene copy number (n = 1080), DNA methylation (n = 345), and phenotypes (n = 1236). In the TCGA database, no significant association was noted between the 11 down-regulated hub genes and patient survival. The findings of ClueGo analysis revealed that three down-regulated KEGG pathways—arachidonic acid metabolism, renin–angiotensin system, and taurine and hypotaurine metabolism—correlated with a decreased survival rate (Figure 4). Based on this finding, we hypothesized that taurine deficiency may lead to increased invasion. In this case, supplement of taurine may reduce invasion of EO771 cells. If this effect is a morphine-opioid receptor-specific action, an antagonist of opioid receptor-naloxone, will not exert similar effect. Moreover, morphine was reported to trigger physiological action through dopamine receptor, so a dopamine receptor antagonist, droperidol, will be tested for its invasion ability. To this end, EO771 cells were treated with morphine (M), morphine+taurine (M+T), naloxone (N), naloxone+taurine (N+T), droperidol (D), droperidol+taurine (D+T) and taurine alone (T) for 4 days to examine their contribution to invasion. In addition, effects of morphine, naloxone, and droperidol on expression of *GAD1* and *CDO1* genes will be determined. As shown in Figure 5, morphine treatment significantly increased invasion compared to no treatment; treatment of taurine alone was not different from no treatment control; and combined treatment of M+T reduced invasion. There was no significant difference among D, N, untreated or taurine groups. No difference was observed between D and D+T, N and N+T. Thus, we conclude that taurine reduced invasion caused by morphine treatment (Figure 5). Based on our RNA-sequencing results, one of taurine biosynthesis enzymes, GAD1, was significantly reduced in tumors derived from morphine-treated mice with lung metastasis (M2) compared to those from saline-treated mice with lung metastasis (S3). Another enzyme, cysteine dioxygenase 1 (CDO1), was not different between these two groups (Figure 5). To determine whether morphine causes the change of these enzymes, EO771 cells treated with morphine, droperidol, or naloxone for 4 days, only morphine treatment reduced expression of *GAD1*. In addition, *CDO1* expression was not affected by morphine, droperidol, or naloxone (Figure 5), consistent with our RNA-sequencing results. *In vivo* results showed a more profound reduction of *GAD1*, possibly due to long-term morphine challenge.

**Figure 4.**
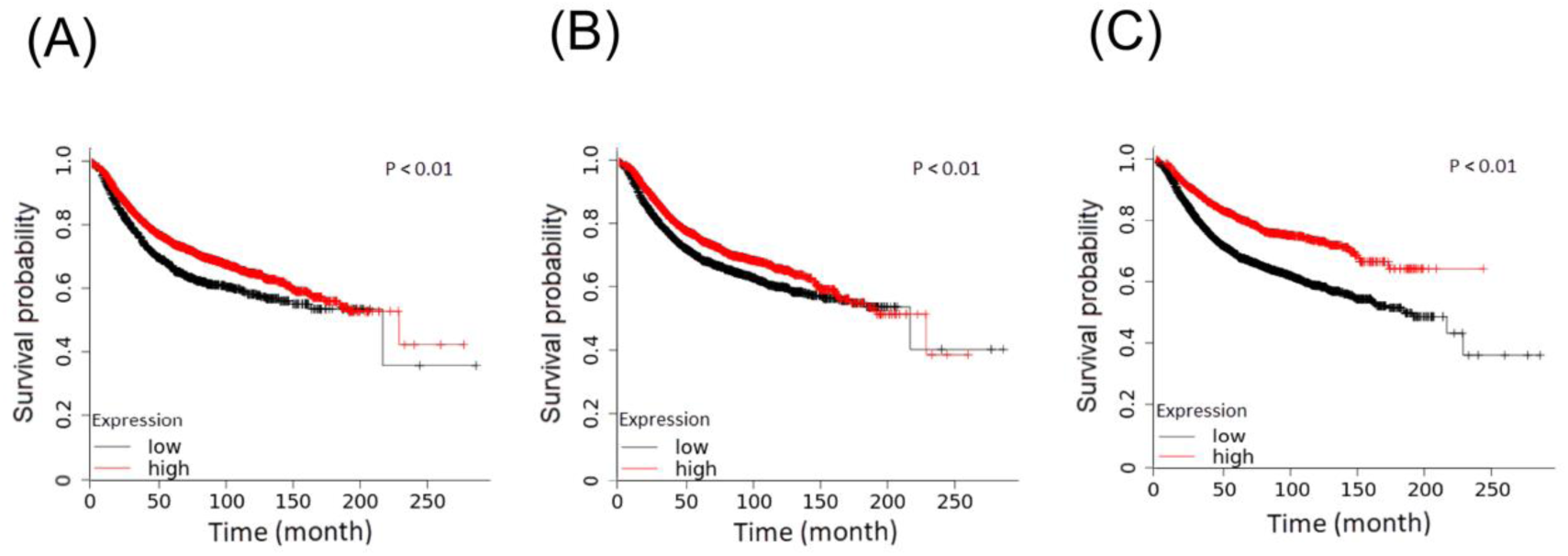
Kaplan–Meier illustration of Kyoto Encyclopedia of Genes and Genomes dominant pathway. Association of high/low expression of genes with survival probability. (A) Kaplan–Meier plot of arachidonic acid metabolism. (B) Kaplan–Meier plot of the renin–angiotensin system. (C) Kaplan–Meier plot of taurine and hypotaurine metabolism. Red line: effect of higher expression of each pathway on survival probability; black line: effect of lower expression of each pathway on survival probability. P < 0.05 indicates a significant difference.

**Figure 5.**
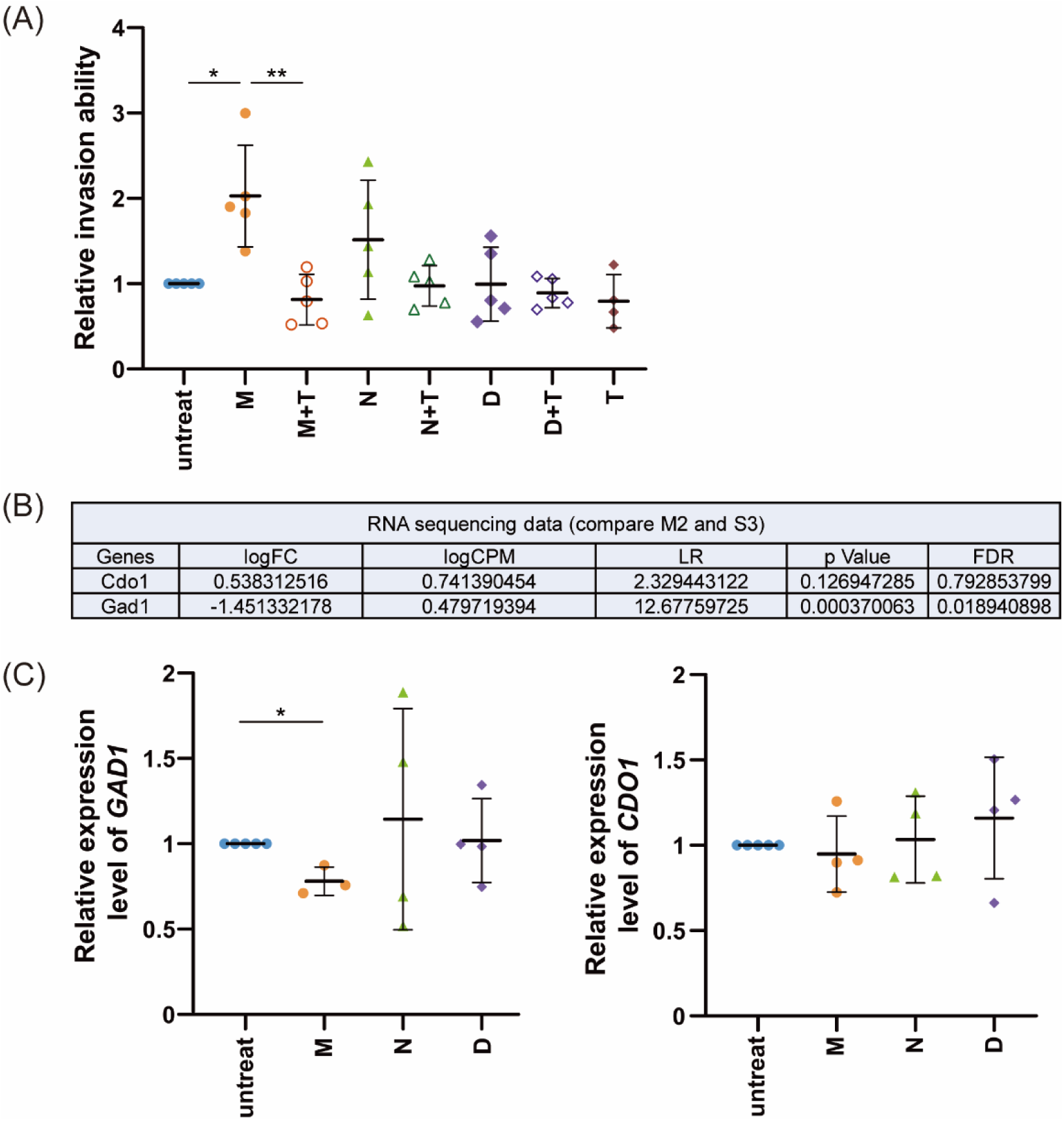
Effect of morphine, naloxone and droperidol treatment on EO771 cell invasion with or without taurine and on expressions of *GAD1* and *CDO1* genes. (A) EO771 cells were treated with 10 μM morphine, 1 μM naloxone (opioid receptor antagonist), 10 μM droperidol (dopamine receptor antagonist) with or without 20 mM taurine every day for 4 days. Boyden chamber assays were then performed to determine relative invasion ability for 26 h. (B) RNA sequencing was performed on tumors derived from saline-treated or morphine-treated mice. S3: saline-treated mice with lung metastasis; M2: morphine-treated mice with lung metastasis (C) EO771 cells were treated with 10 μM morphine, 1 μM naloxone or 10 μM droperidol every day for 4 days. RNAs of EO771 cells were collected. Relative expression levels of *GAD1* and *CDO1* genes were determined via qPCR, normalized to *GAPDH* level and no treatment group. M: morphine; N: naloxone; D: droperidol; T: taurine. (* indicates a significant difference of P < 0.05 between groups; ** indicates a significant difference of P < 0.01 between groups, determined using the Student’s t test.). Data were from at least three independent experiments.

## Discussion

Whether morphine promotes or inhibits tumor formation remains controversial [28] possibly due to the various types of tumors and durations of morphine usage. Although morphine has been implicated in immune modulation [29, 30], this role has not been linked to tumor formation or invasion. We identified in this study the majority of morphine-mediated pathways, especially taurine/hypotaurine metabolism (mmu00430) in DAVID and ClueGo analysis. Further analysis showed that long-term exposure correlated with lower *GAD1* and *CDO1* levels. In TCGA database, human triple negative breast cancer showed decreasing *GAD1* and *CDO1* level at late stage (Figure S4). Our findings suggest that long-term morphine usage promotes the metastasis of triple-negative breast cancer, likely through reducing the expression of taurine biosynthesis enzymes.

Taurine is an abundant free amino acid which is involved in anti-inflammation [31], improves the effect of antioxidants, enhances immune response to against cancer cell, and induces cancer cell apoptosis [32, 33]. Taurine exerts both suppressive and cytotoxic effects on tumors [31]. Relative taurine levels have been implicated as diagnostic markers for tumors [34]. Moreover, taurine attenuates tumor growth and invasion by suppressing extracellular-signal-regulated kinase/ribosomal S6 kinase signaling [31, 35]. Thus, decreased taurine levels may be associated with poor prognosis [31]. The *CDO1* and *GAD1* genes encode key enzymes during taurine biosynthesis [36]. Thus, decreased expression of these two genes leads to decreased taurine synthesis. The complexity of morphine-mediated effects is partly dependent on the exposure duration. Short-term morphine exposure leads to an increased *GAD1* level [37], whereas long-term morphine exposure results in a decreased *GAD1* level [38, 39]. Consistent with this finding, our results indicated that morphine usage led to decreased *GAD1* expression. *CDO1* is a metastasis-related gene, and its suppression is associated with poor prognosis [40]. In our study, lower expression level of *GAD1* caused by long-term morphine usage may have been the key mechanism underlying the reduction in taurine level and thus the promotion of tumor invasion (Figure 6). Silencing of the *GAD1* and *CDO1* genes is involved in the poor prognosis of patients with renal cell carcinoma [41]. This result suggests that a lower taurine level is essential in the advanced stages of cancers. Human triple-negative breast cancer datasets revealed decreased *CDO1* expression during cancer progression, whereas *GAD1* expression initially increased and then decreased in stage IV (Figure S4). This finding strongly suggests a molecular change during cancer advancement. Long-term morphine usage promotes the metastasis of triple-negative breast cancer, likely through reducing the expression of taurine biosynthesis enzymes.

**Figure 6.**
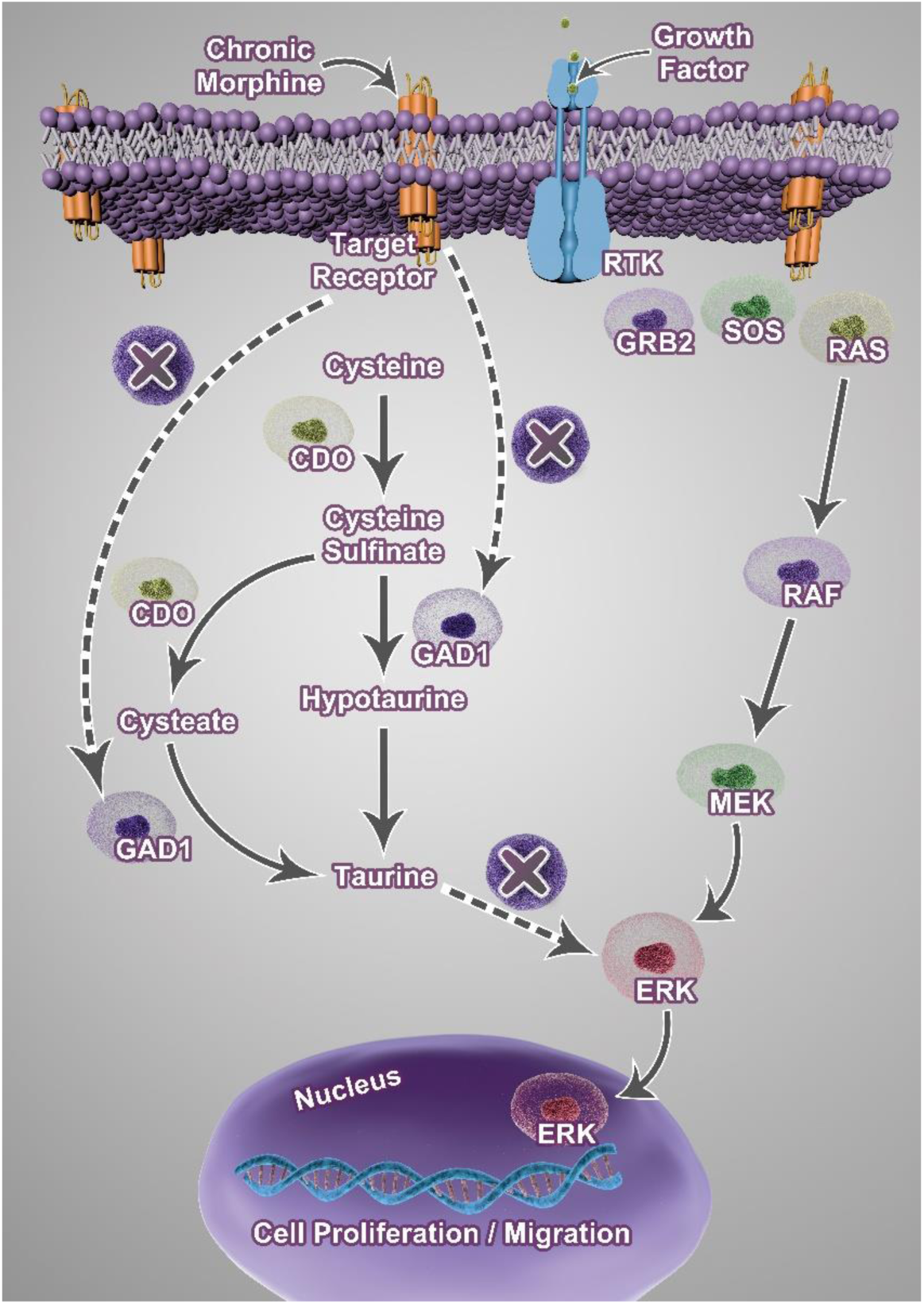
Illustration of morphine-mediated suppression of taurine level. Morphine acts through candidate target receptors (μ opioid receptor and/or dopamine receptor) and reduces expression of *GAD1*, whose activity is responsible for biosynthesis of taurine. Taurine attenuates tumor progression by inhibiting the ERK pathway. *GAD1*: glutamate decarboxylase 1; *CDO1*: cysteine dioxygenase; RTK: receptor of tyrosine kinase.

In our study, we found that the renin–angiotensin system is down-regulated in triple-negative breast cancer cells after long-term morphine exposure. Inhibition of the renin–angiotensin system can be beneficial for cancer treatment [42]. Angiotensin-converting enzyme II gene expression inhibits neoangiogenesis [43], and its decreased expression may lead to poor survival. In our TCGA breast cancer database analysis, inhibition of the renin–angiotensin system was correlated with poor survival. A previous study reported that the arachidonic acid level may not exert a causal effect on cancer growth. Other metabolites involved in the arachidonic acid pathway, such as 20-hydroxy-eicosatetraenoic acid (*HETE*), may promote tumor growth and invasion [44, 45]. In our study, we noted the down regulation of genes involved in arachidonate metabolism, including *PTGS2*, epoxide hydrolase 2, and phospholipase A2. Long-term morphine usage is related to the inhibition of substance P [46], and substance P induces the arachidonate cascade by stimulating 12-HETE [47]. Therefore, the definitive correlation between tumor metastasis and arachidonate metabolism will need to be clarified in future studies.

Our study has some limitations. To mimic long-term morphine usage, mice were administered morphine 2 weeks before EO771 cell implant. This is due to the fast tumor-forming property of EO771 cells *in vivo*. Thus, the experimental design might not perfectly recapitulate the clinical setting of long-term morphine usage for pain relief. Nonetheless, using mouse-origin of EO771 cells allows us to test the effect of morphine in normal mice instead of using nude mice (immune-deficient mice). This is crucial in our study design based on the known immune-suppressing property of morphine. In addition, our metastasis tumor samples did not have same-day controls. When primary tumors were removed from mice fat pad, the tissue has been destroyed due to large tumors. Thus, it is not possible to obtain same-day, non-metastasis tumors as control. To circumvent this problem, the *in vitro* cell experiments were designed to test whether the identified down-regulation of *GAD1* gene was indeed morphine-dependent, as proof-of-concept experiments. Moreover, our *in vivo* and *in vitro* experiments did not echo each other perfectly on morphine-induced tumor growth. Based on the *in vitro* data, morphine promoted tumor cell invasion without increased cell proliferation (Figure 1), whereas morphine increased tumor sizes by Day 23 in animal experiments (Figure 2). We think morphine-promoted tumor growth likely results from immune suppression and/or tumor microenvironmental change, such as metabolism, *in vivo*. In the *in vitro* tumor cell only model, one cannot recapitulate this effect.

In conclusion, this study reveals one of the mechanisms that underlies morphine-promoted metastasis of triple negative breast cancer through reducing the level of taurine biosynthesis enzyme.

## Abbreviation

GPCR: G-protein-coupled receptor
MOR: μ-opioid receptor
NK: natural killer
FBS: fetal bovine serum
MTT: 3-(4,5 dimethylthiazol-2-tl)-2,3-diphenyltetrazolium bromide
BSA: Bovine serum albumin
NTHU: National Tsing Hua University
GAPDH: Glyceraldehyde-3-phosphate dehydrogenase
KEGG: Kyoto Encyclopedia of Genes and Genomes
GO: Gene Ontology
DEGs: Differentially expressed genes
DAVID: Database for Annotation, Visualization, and Integrated Discovery
PCA: Principal component analysis
PPIs: Protein–protein interactions
TCGA: Cancer Genome Atlas
GAD1: Glutamate decarboxylase 1
BP: Biological process
MF: Molecular function
CC: Cellular compartment
TTN: titin
TCAP: titin cap
ACTA1: actin alpha 1, skeletal muscle
MYH1: myosin heavy chain 1
TNNT3: troponin T3,
MYL1: myosin light chain 1
MYH4: myosin heavy chain 4
TNNI2: troponin I2
ATP2a1: atpase sarcoplasmic/endoplasmic reticulum Ca2+ transporting 1
TNNC2: troponin C2
CDO1: Cysteine dioxygenase 1
SREBF1: Sterol regulatory element-binding transcription factor 1
TCF3: Transcription factor 3
PTGS2: Prostaglandin-endoperoxide synthase 2
HETE: 20-hydroxy-eicosatetraenoic acid
HNF1A-AS1: Epatocyte nuclear factor 1 homeobox A-antisense RNA 1

## Acknowledgements

This work is supported by the Ministry of Science and Technology, Taiwan (grant#: 108-2320-B-007-005-MY3 and 111-2311-B-007-012 to Dr. Linyi Chen) and National Tsing Hua University, Taiwan (grant#: 110F7MBNLE1 to Dr. Shih-Hong Chen). This manuscript was edited by Wallace Academic Editing

## Declaration

The authors declare that they have no known competing financial interests or personal relationships that could have appeared to influence the work reported in this paper.

All authors did not use AI-assisted -technology.

## Author contribution and contributor statement

**Shih-Hong Chen**: Data analysis, Investigation, Visualization, Writing – original draft, review & editing, Funding. **Chien-Hung Shih**: Animal study, quantification of tumor area in lung tissue, Writing –original draft. **Ting-Ling Ke**: Conducted all experiments for revision. Data analysis, Writing – review and editing. **Zi-Xuan Huang**: Tumor RNA preparation. **Kuo-Chin Chen**: TCGA datasets analysis, Software, Visualization. **Chia-Ni Hsiung**: Software, Visualization, Statistical analysis. **Tsung-Hsien Chuang**: Cell line, Methodology. **Li-Kuei Chen**: Conceptualization, Provide reagents, Funding. **Linyi Chen**: Conceptualization, Investigation, Visualization, Writing –original draft, review & editing, Funding.

**Linyi Chen** is responsible for the overall content as guarantor, accepts full responsibility for the finished work, the conduct of the study, had access to the data, and controlled the decision to publish.

## Availability of data

The datasets generated and/or analyzed during the current study are available in the Figshare repository, DOI 10.6084/m9.figshare.22644256

## Ethics approval and consent to participate

Animal usage protocols were reviewed and approved by NTHU’s Institutional Animal Care and Use Committee (approval number: 10607).

## Consent for publication

Not applicable

---

**Figure S1.**
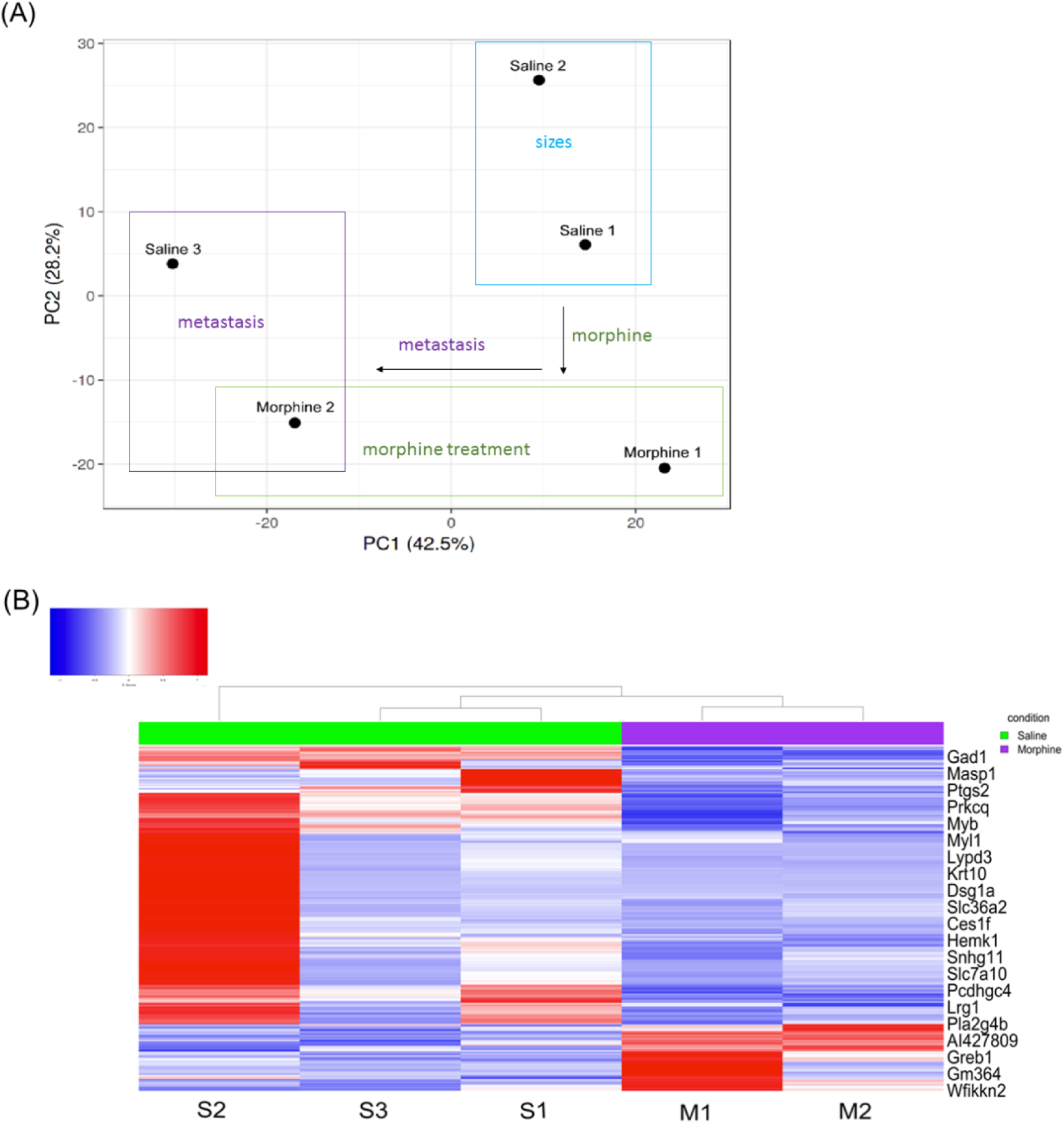
Principle component analysis and heatmap illustration. (A) Principal component analysis. Saline 1 group (S1) had tumor size < 50% mean volume without metastasis, Saline 2 group (S2) had tumor size > 50% mean volume without metastasis, Saline 3 group (S3) had tumor metastasis, Morphine 1 (M1) group did not follow metastasis, and Morphine 2 group (M2, Morphine+metastasis) had tumor metastasis. (B) Heatmap of all five groups.

**Figure S2.**
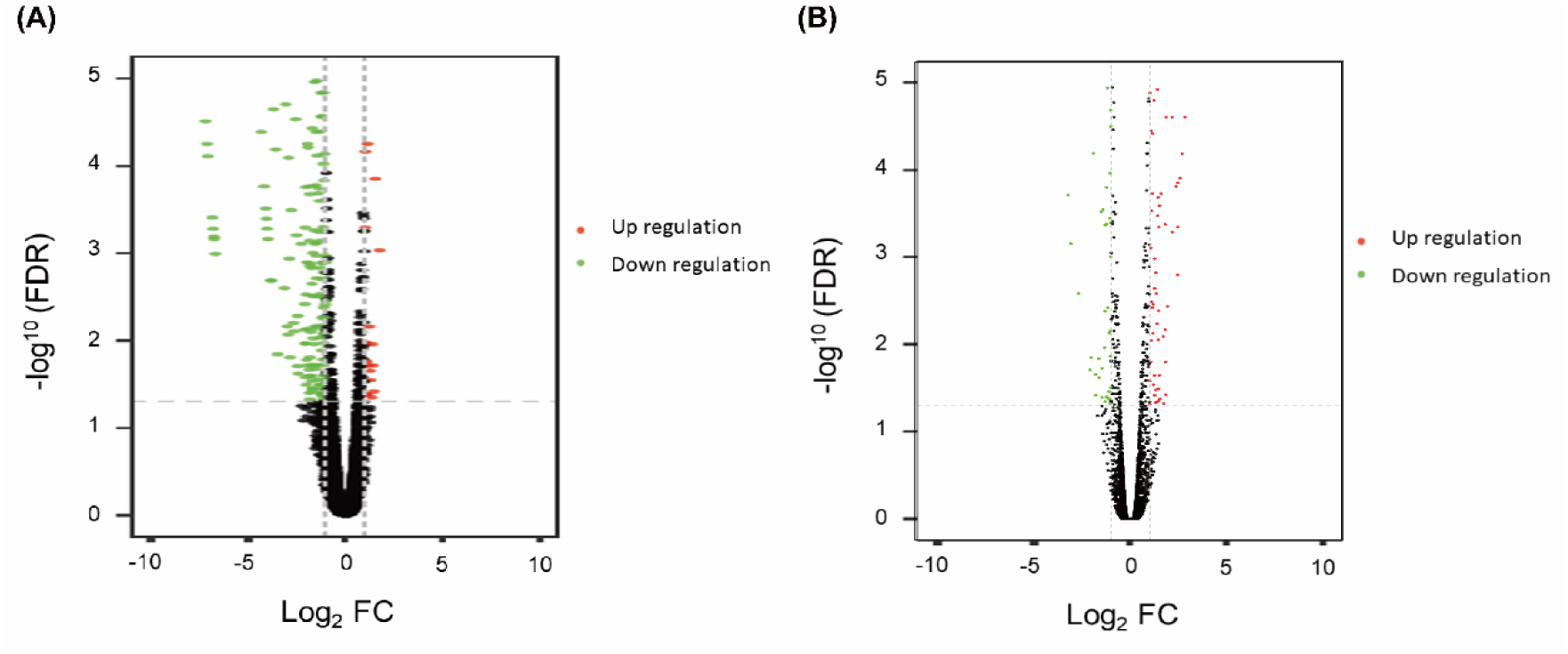
Volcano plot of up-regulated and down-regulated genes. Up-regulated genes are designated in red, whereas down-regulated genes are in green. Red spot: dominant up-regulated expression (fold-change > 1. FDR < 0.05); green spot: dominant down-regulated expression (fold-change < −1, FDR <0.05); black spot: non-dominant fold-change. (A) Compare M2 and the average of S1 and S2 groups. (B) Compare M2 and S3 groups.

**Figure S3.**
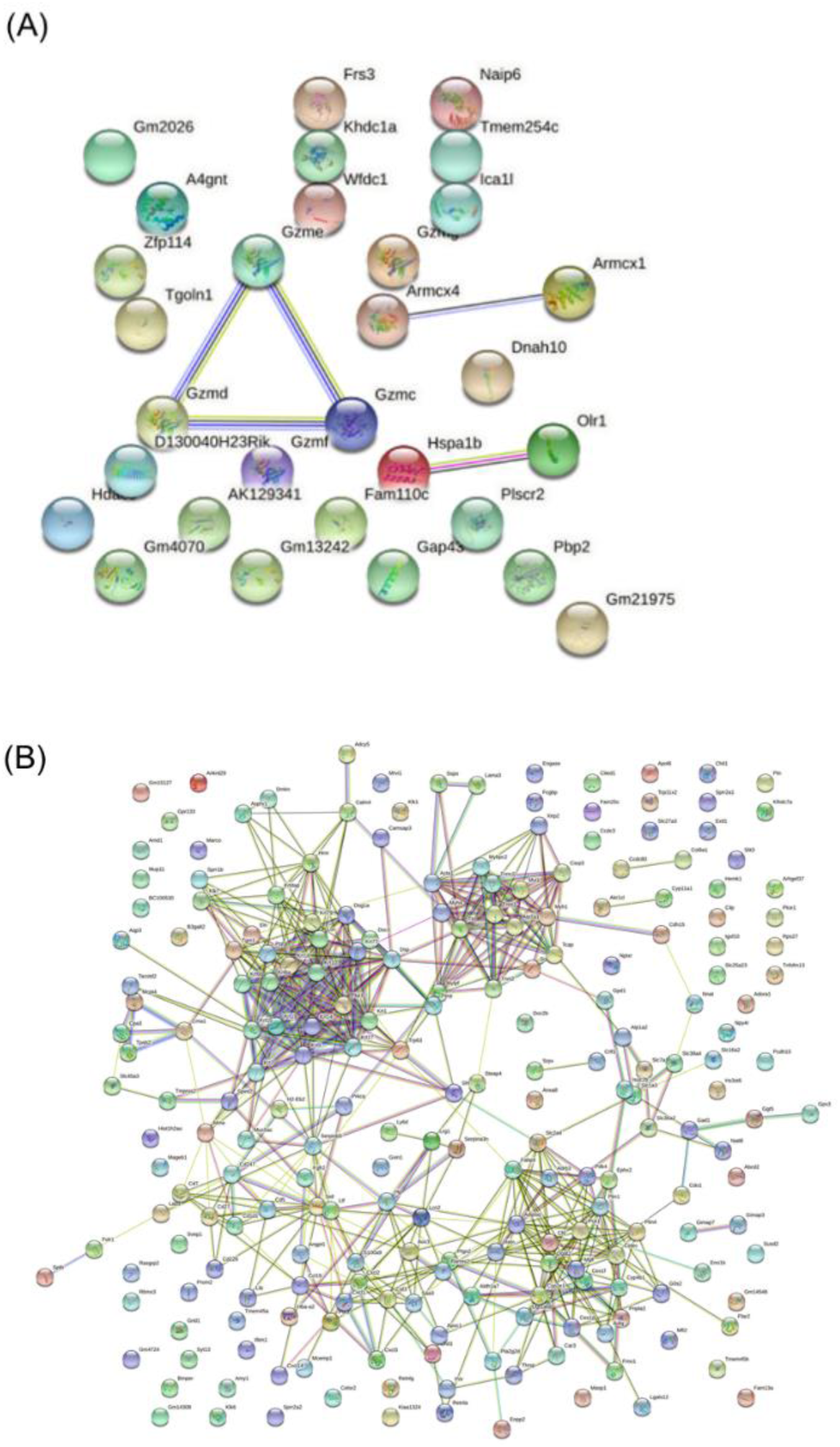
Predicted protein–protein interaction (PPI) by using STRING. (A) PPI according to the dominant up-regulated genes (fold-change > 1, FDR < 0.05). (B) PPI based on dominant down-regulated genes (fold-change < −1, FDR < 0.05). Colored nodes: first level interactors of query proteins; white nodes: second-level interactors. Green-blue line: curated databases; purple line: experimentally determined; green line: predicted gene neighborhood; red line: predicted gene fusions; blue line: predicted gene co-occurrence; light-green line: text mining; black line: co-expression; light-blue line: protein homology

**Figure S4.**
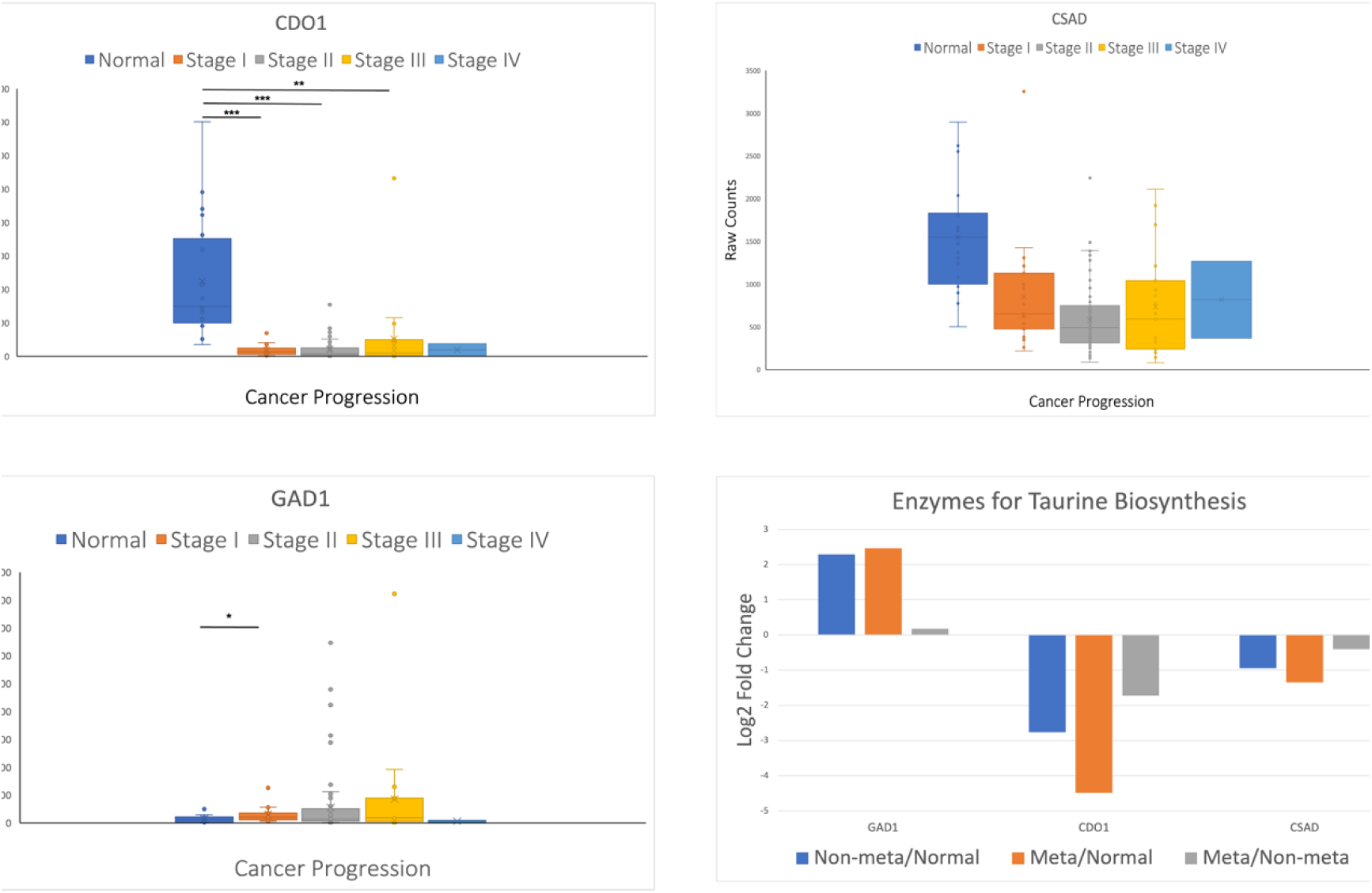
Expression of taurine biosynthesis genes in TCGA database. Fold change of gene expression for taurine biosynthesis genes, including *GAD1, CDO1*, and *CSAD*. Raw counts of *GAD1*, *CDO1*, and *CSAD* were compared in groups of normal tissue and stage I-IV. Relative non-meta/normal, meta/normal, and meta/non-meta level of *GAD1*, *CDO1*, and *CSAD* expression were shown. For *GAD1*, non-meta/normal: P < 0.01, meta/normal: P = 0.04, and meta/non-meta: P = 0.99. For *CDO1*, non-meta/normal: P < 0.01, meta/normal: P < 0.01, and meta/non-meta: P = 0.57. For *CSAD*, non-meta/normal: P < 0.01, meta/normal: P = 0.02, and meta/non-meta: P = 0.94. Cancer patients: 116 people, Normal: 20 people.

**Supplementary Table 1.** Grouping of tumors based on sizes, gene expression, metastasis and treatment.

| Category<br>(Submit for RNAseq) |  | S1 | S2 | S3 | M1 | M2 |  |
| --- | --- | --- | --- | --- | --- | --- | --- |
| Isolated tumors |  | s2-1+s1 | s4-1+s2+s3 | s4+s5 | m4-1+m3 | m1+m6 |  |
| Based on |  | Gene expression | Tumor small | Tumor large +Metastasis | Gene expression | Metastasis |  |
|  |  | Relative gene expression |  |  |  |  |  |
| Category | Tumors | EGFR | HER2 | GLI1 | EZH2 | Tumor volume (mm<sup>3</sup>) | Based on |
| S1 | saline 2-1 | 2.98 | 1.88 | 4.15 | 1.12 | 125 | Gene expression |
|  | saline 1 | 3.67 | 1.67 | 4.93 | 1.34 | 114 |  |
| S2 | saline 4-1 | 3.53 | 1.57 | 5.22 | 0.79 | 4 | Small size tumors |
|  | saline 2 | 3.82 | 1.54 | 4.85 | 1.12 | 75 |  |
|  | saline 3 | 1.50 | 1.33 | 4.75 | 1.13 | 52 |  |
| S3 | saline 4 | 1.85 | 1.63 | 4.76 | 1.02 | 501 | Large size tumors +Metastasis |
|  | saline 5 | 2.55 | 1.12 | 3.09 | 0.91 | 325 |  |
|  |  | Relative gene expression |  |  |  |  |  |
| Category | Tumors | HER2 | GLI1 | EZH2 | Tumor volume (mm<sup>3</sup>) | Based on |  |
| M1 | morphine 4-1 | 2.51 | 8.12 | 1.76 | 574 | Gene expression |  |
|  | morphine 3 | 2.59 | 5.30 | 1.78 | 645 |  |  |
| M2 | morphine 1 | 2.17 | 10.37 | 1.90 | 685 | Metastasis |  |
|  | morphine 6 | 2.88 | 20.22 | 1.73 | 549 |  |  |

**Supplement Table 2.**
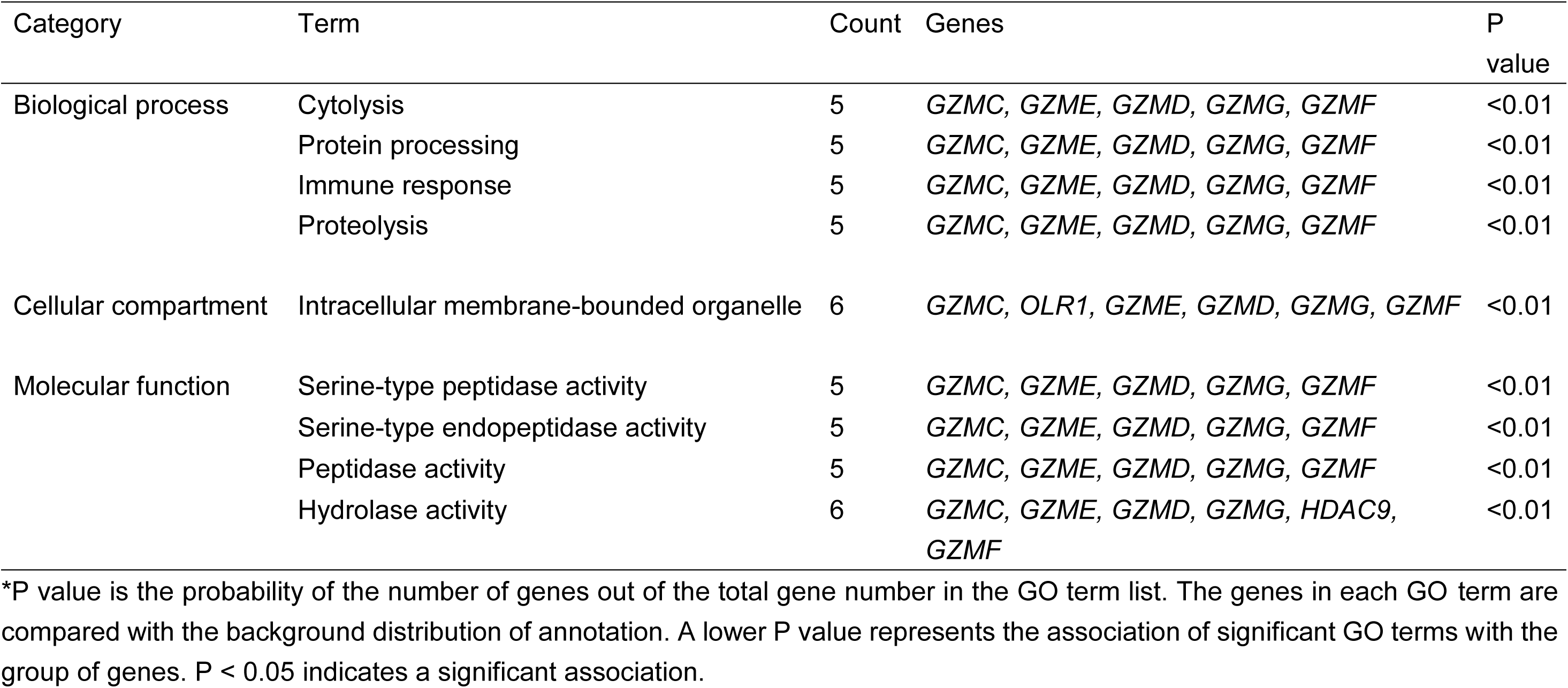
Gene ontology analysis of upregulated differentially expressed genes.

| Category | Term | Count | Genes | P value |
| --- | --- | --- | --- | --- |
| Biological process | Cytolysis | 5 | *GZMC, GZME, GZMD, GZMG, GZMF* | <0.01 |
|  | Protein processing | 5 | *GZMC, GZME, GZMD, GZMG, GZMF* | <0.01 |
|  | Immune response | 5 | *GZMC, GZME, GZMD, GZMG, GZMF* | <0.01 |
|  | Proteolysis | 5 | *GZMC, GZME, GZMD, GZMG, GZMF* | <0.01 |
| Cellular compartment | Intracellular membrane-bounded organelle | 6 | *GZMC, OLR1, GZME, GZMD, GZMG, GZMF* | <0.01 |
| Molecular function | Serine-type peptidase activity | 5 | *GZMC, GZME, GZMD, GZMG, GZMF* | <0.01 |
|  | Serine-type endopeptidase activity | 5 | *GZMC, GZME, GZMD, GZMG, GZMF* | <0.01 |
|  | Peptidase activity | 5 | *GZMC, GZME, GZMD, GZMG, GZMF* | <0.01 |
|  | Hydrolase activity | 6 | *GZMC, GZME, GZMD, GZMG, HDAC9, GZMF* | <0.01 |

\*P value is the probability of the number of genes out of the total gene number in the GO term list. The genes in each GO term are compared with the background distribution of annotation. A lower P value represents the association of significant GO terms with the group of genes. $P < 0.05$ indicates a significant association.

**Supplementary Table 3.**
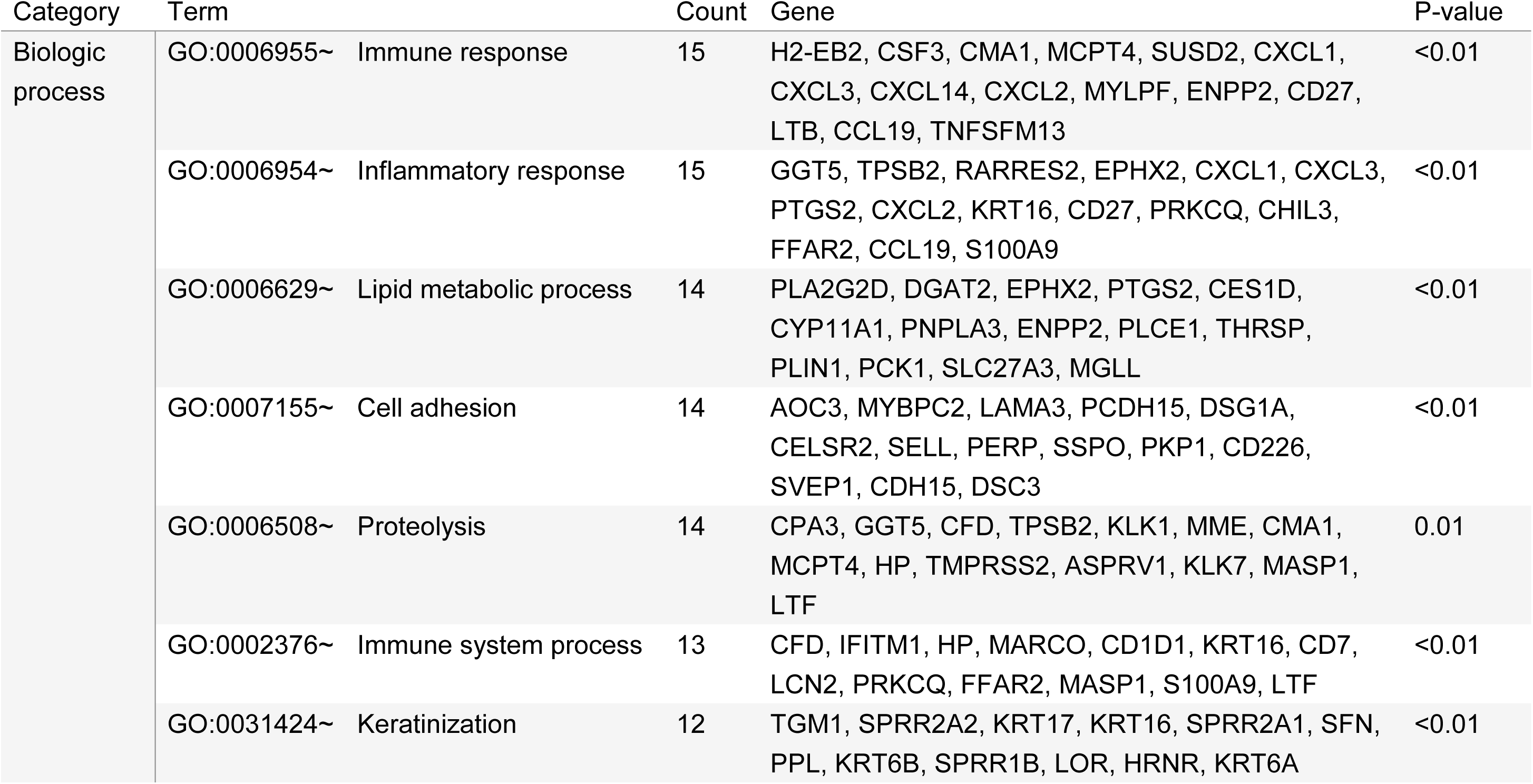
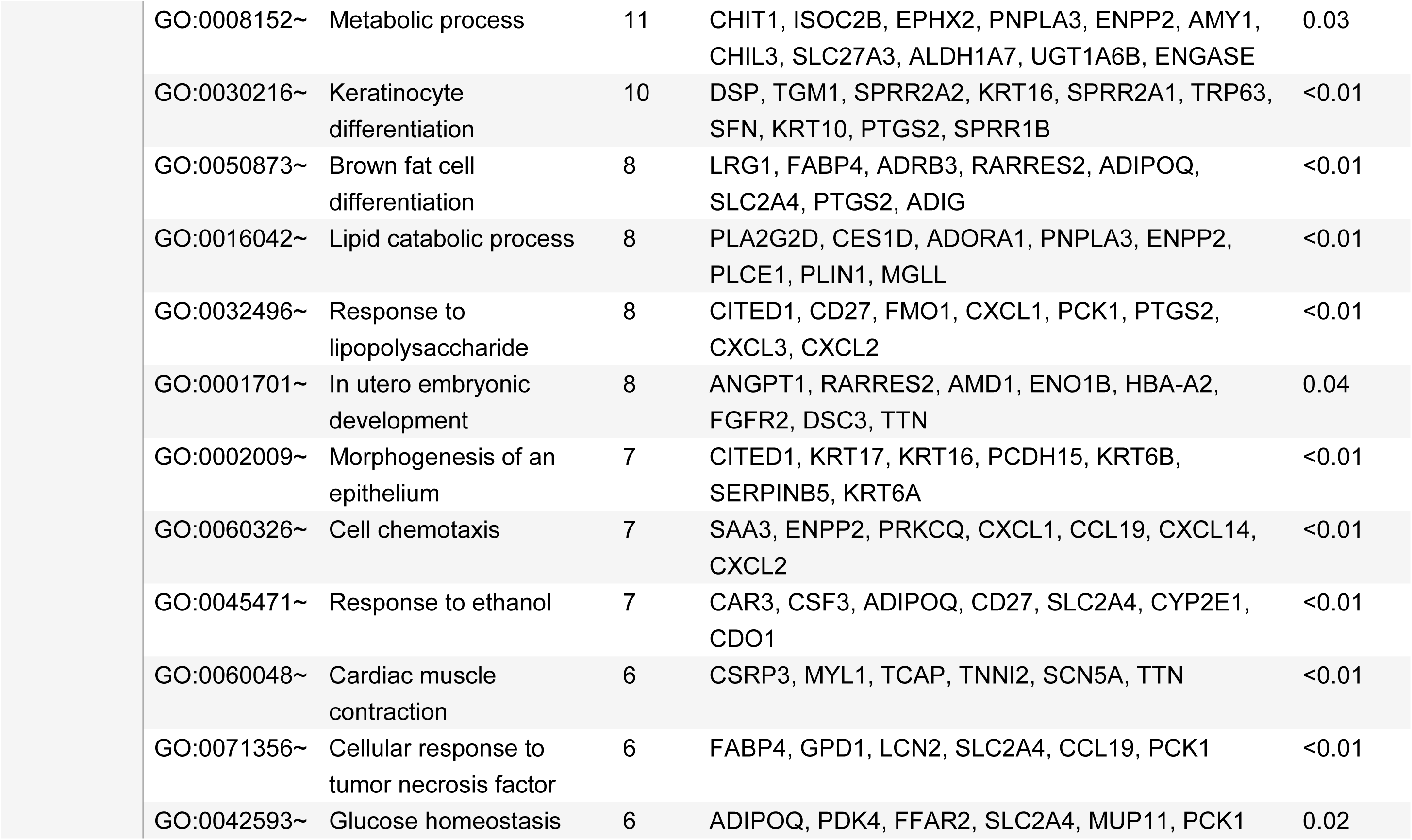
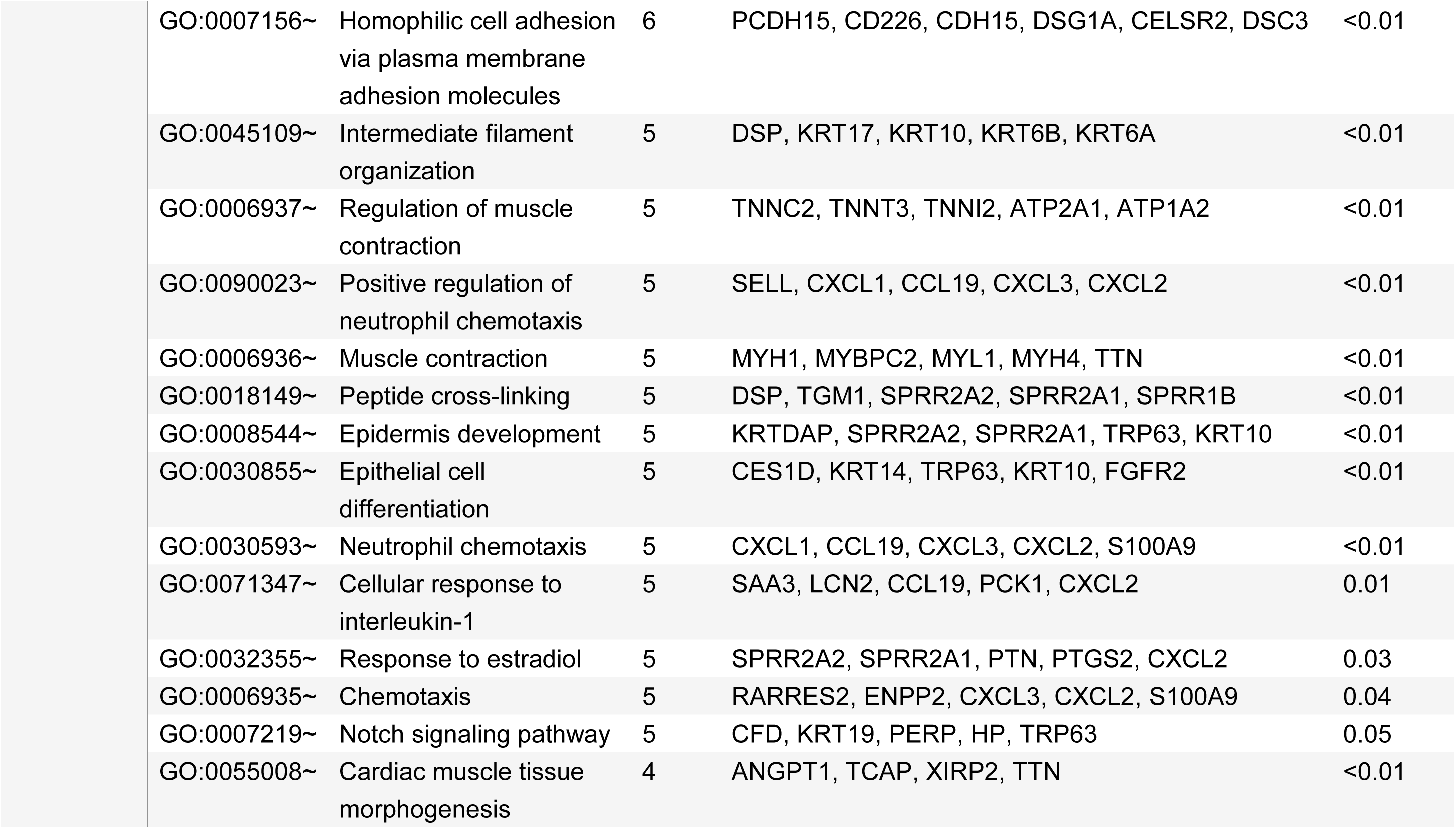
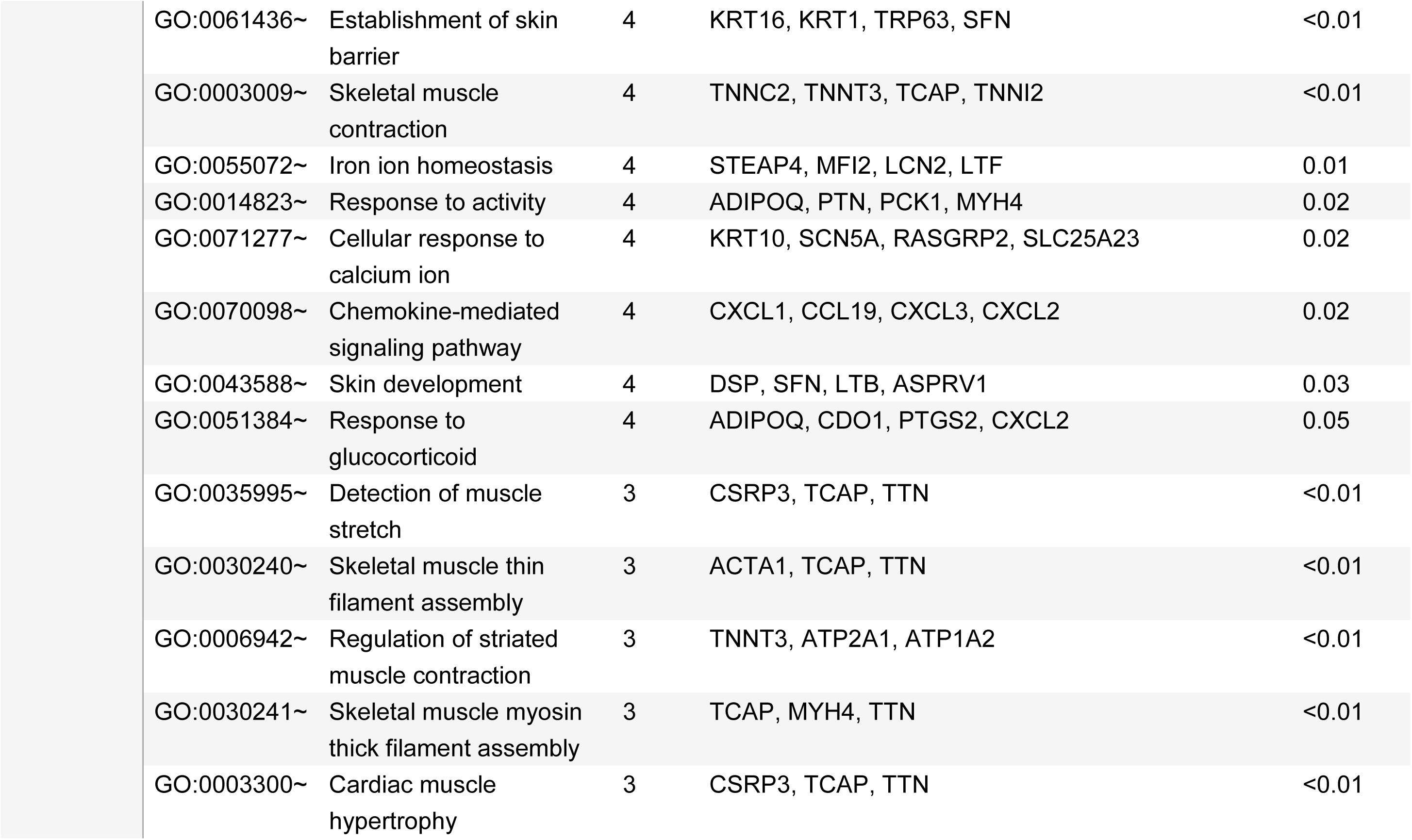
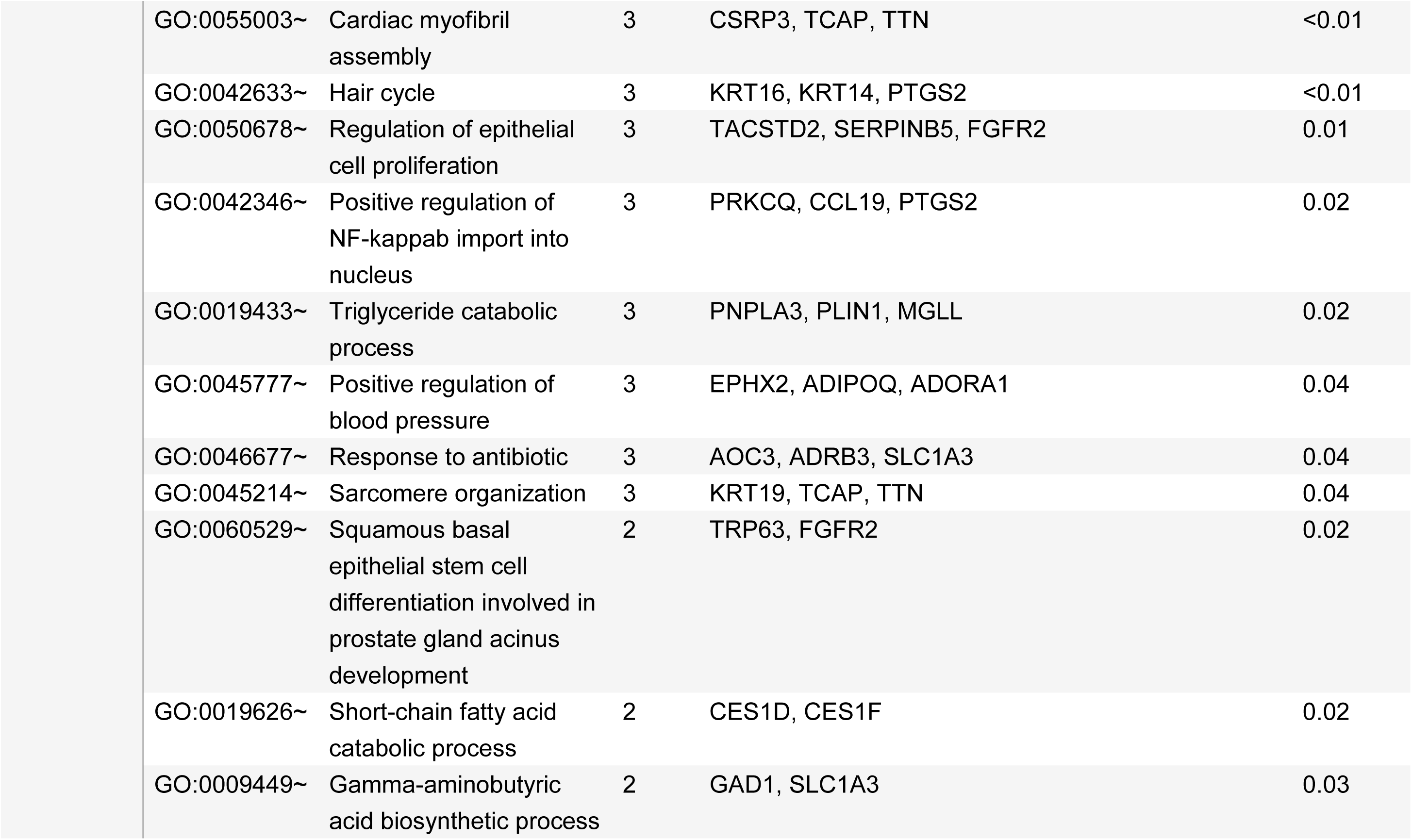
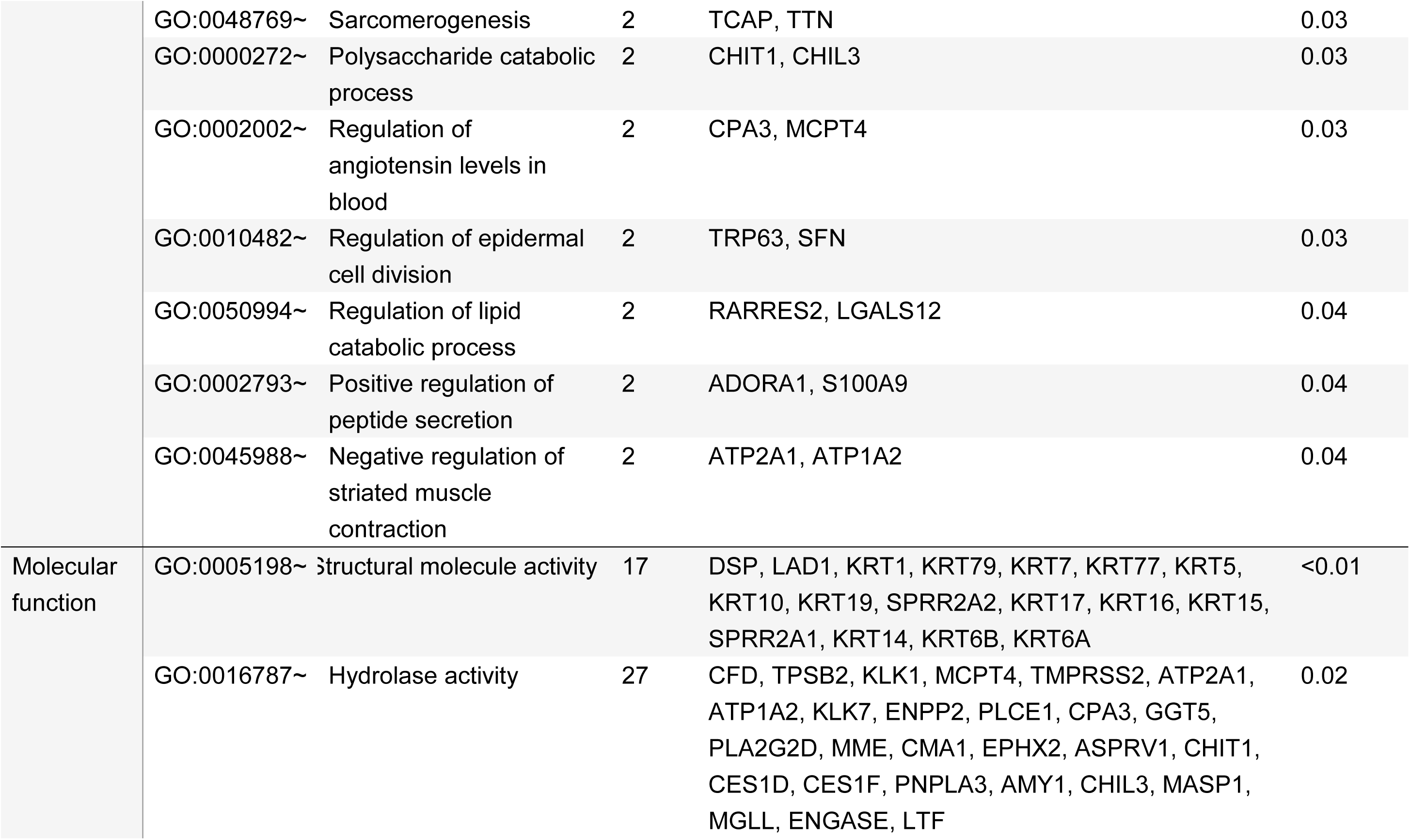
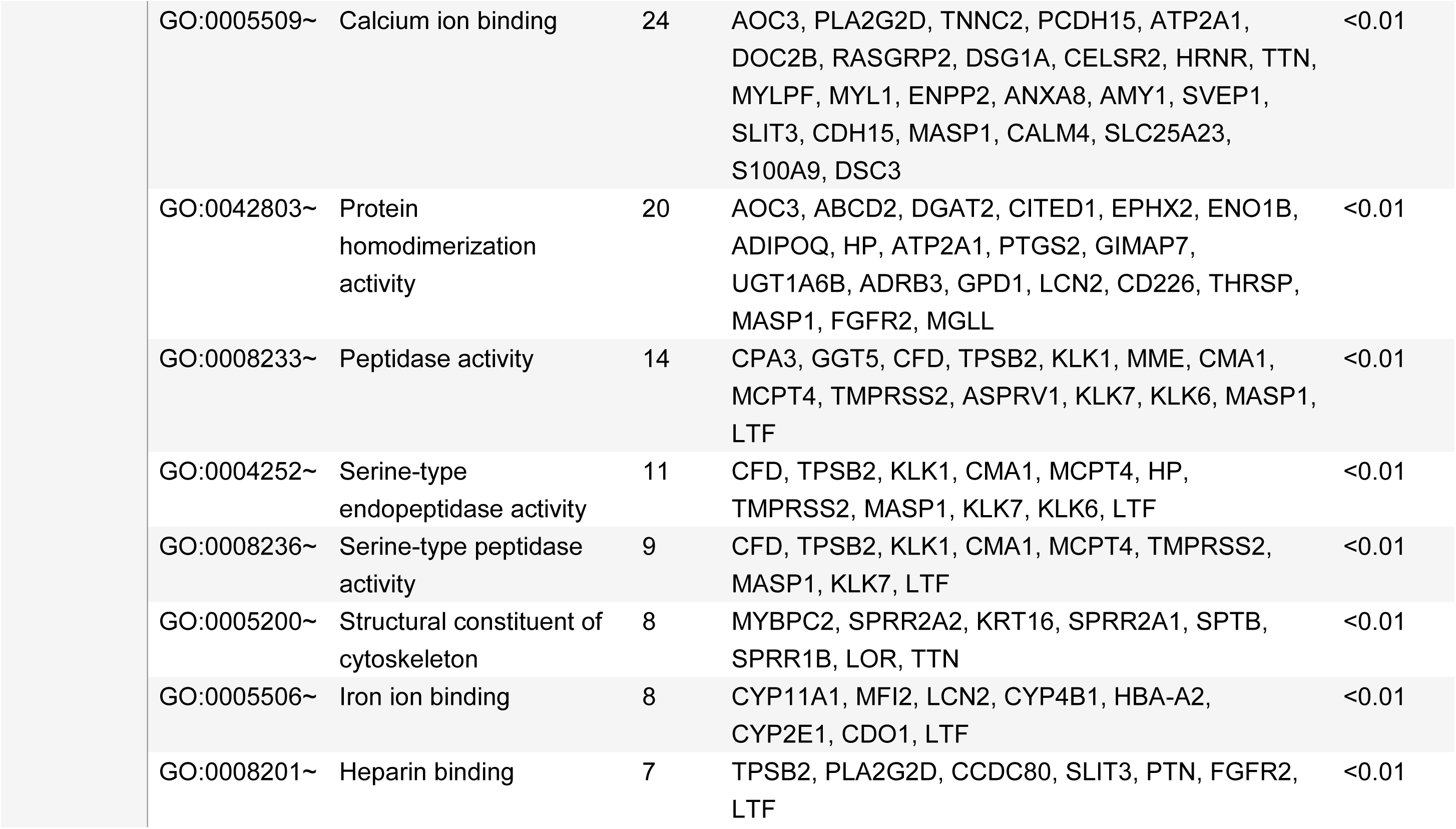
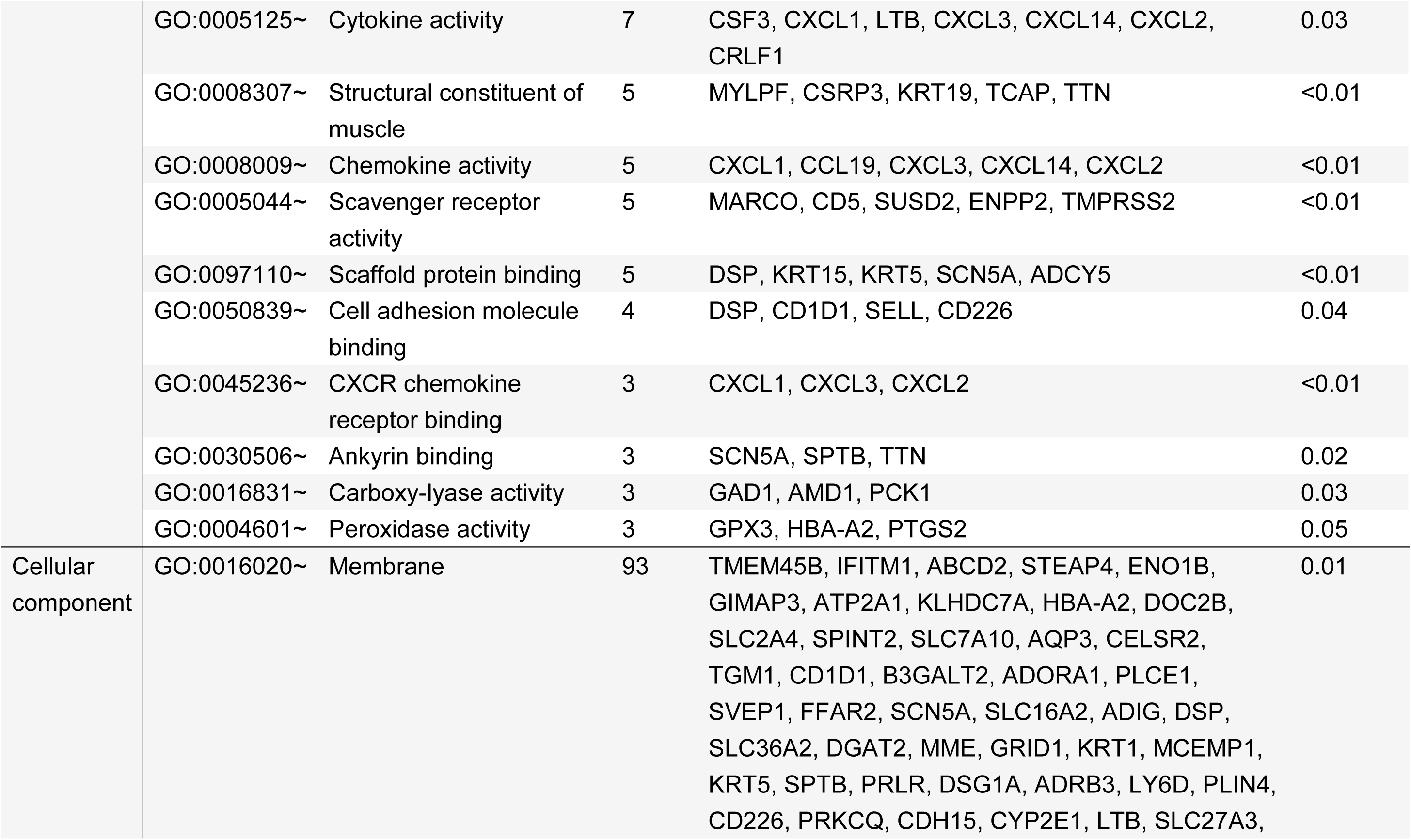
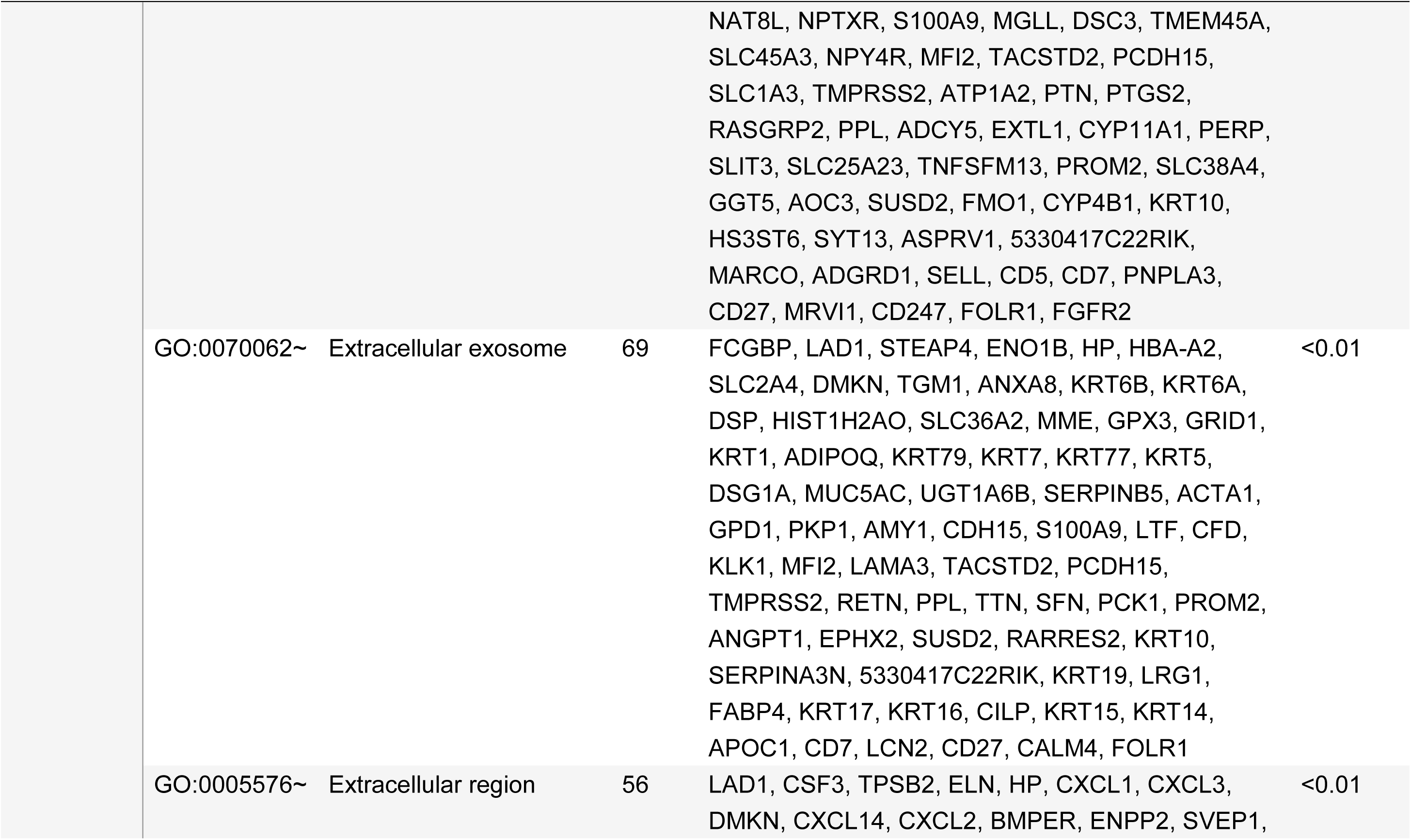
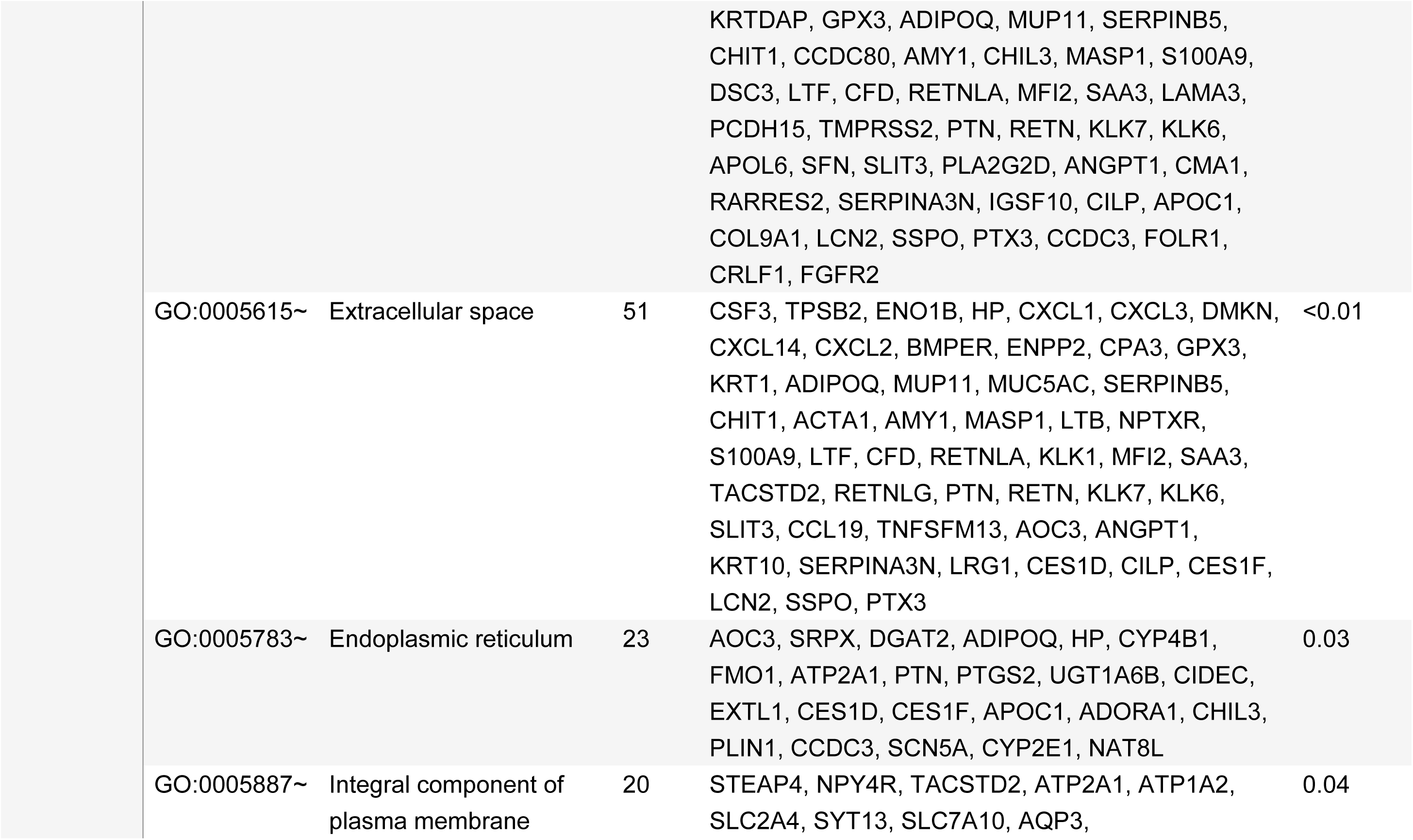
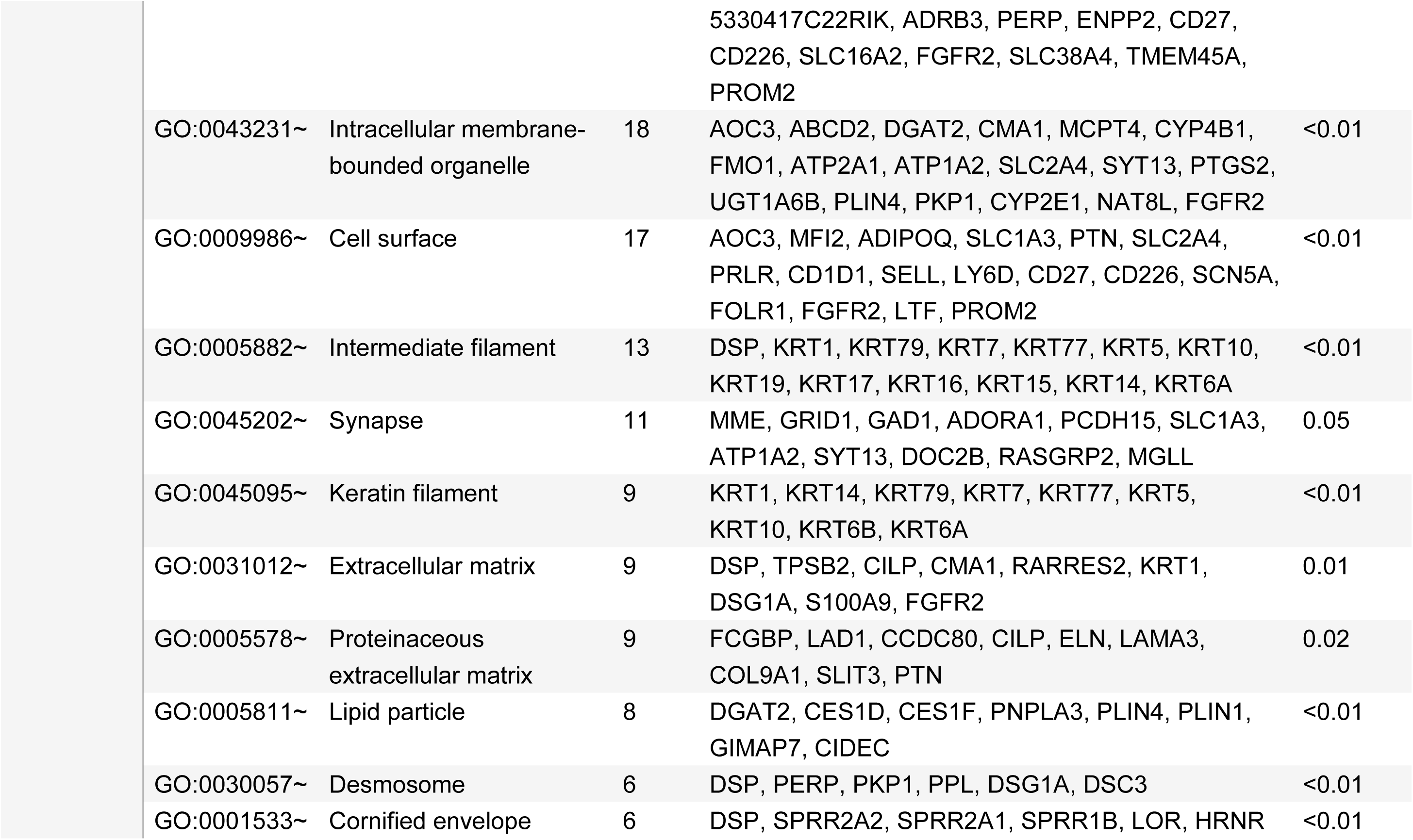
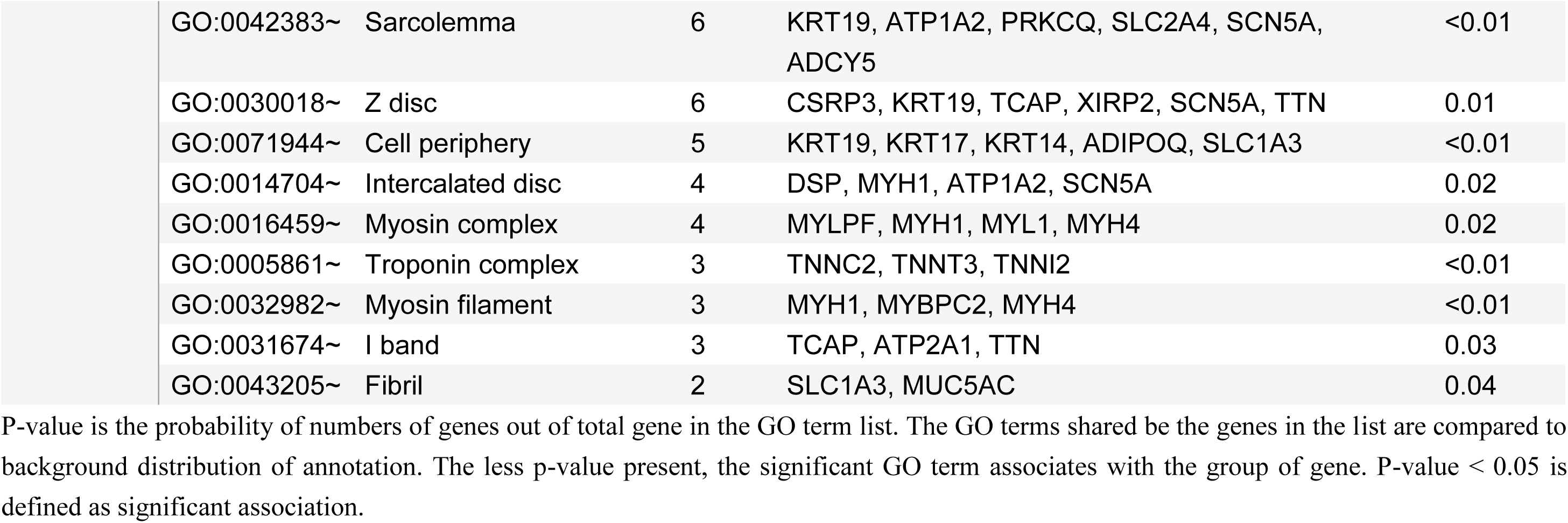
Morphine-effect of downregulation genes for Gene Ontology terms.

**Supplementary Table 4.** KEGG analysis.

| Pathway name | Count | Genes | P value |
| --- | --- | --- | --- |
| **Upregulation** |  |  |  |
| Legionellosis | 2 | *NAIP7, HSPA1B* | 0.03 |
| **Downregulation** |  |  |  |
| Cytokine–cytokine receptor interaction | 8 | *CSF3, CD27, CXCL1, LTB, CCL19, CXCL14, CXCL2, PRLR* | 0.033 |
| Regulation of lipolysis in adipocytes | 7 | *FABP4, ADRB3, ADORA1, PLIN1, PTGS2, MGLL, ADCY5* | <0.01 |
| cGMP-PKG signaling pathway | 7 | *ADRB3, ADORA1, ATP2A1, MRVI1, ATP1A2, CALM4, ADCY5* | 0.02 |
| Chemokine signaling pathway | 7 | *CXCL1, CCL19, RASGRP2, CXCL3, CXCL14, CXCL2, ADCY5* | 0.04 |
| Arachidonic acid metabolism | 6 | *GGT5, PLA2G2D, GPX3, EPHX2, CYP2E1, PTGS2* | <0.01 |
| Hematopoietic cell lineage | 5 | *CSF3, CD1D1, MME, CD5, CD7* | 0.02 |
| Protein digestion and absorption | 5 | *CPA3, MME, ELN, COL9A1, ATP1A2* | 0.02 |
| Pancreatic secretion | 5 | *CPA3, PLA2G2D, ATP2A1, ATP1A2, ADCY5* | 0.04 |
| Renin–angiotensin system | 4 | *CPA3, KLK1, MME, CMA1* | <0.01 |
| Taurine and hypotaurine metabolism | 3 | *GGT5, GAD1, CDO1* | <0.01 |

\*P value is the probability of the number of genes out of the total gene number in the KEGG list. The genes in the KEGG list are compared with the background distribution of annotation. A lower P value represents the association of significant KEGG pathways with the group of genes. $P < 0.05$ indicates a significant association

